# Cortical glutamatergic projection neuron types contribute to distinct functional subnetworks

**DOI:** 10.1101/2021.12.30.474537

**Authors:** Hemanth Mohan, Xu An, X. Hermione Xu, Hideki Kondo, Shengli Zhao, Katherine S. Matho, Simon Musall, Partha Mitra, Z. Josh Huang

## Abstract

The cellular basis of cerebral cortex functional architecture remains not well understood. A major challenge is to monitor and decipher neural network dynamics across broad cortical areas yet with projection neuron (PN)-type resolution in real time during behavior. Combining genetic targeting and wide-field imaging, we monitored activity dynamics of subcortical-projecting (PT^Fezf2^) and intratelencephalic-projecting (IT^PlxnD1^) types across dorsal cortex of mice during different brain states and behaviors. IT^PlxnD1^ and PT^Fezf2^ neurons showed distinct activation patterns during wakeful resting, spontaneous movements, and upon sensory stimulation. Distinct IT^PlxnD1^ and PT^Fezf2^ subnetworks were dynamically tuned to different sensorimotor components of a naturalistic feeding behavior, and optogenetic inhibition of ITs^PlxnD1^ and PTs^Fezf2^ in subnetwork nodes disrupted distinct components of this behavior. Lastly, IT^PlxnD1^ and PT^Fezf2^ projection patterns are consistent with their subnetwork activation patterns. Our results show that, in addition to the concept of columnar organization, dynamic areal and PN type-specific subnetworks are a key feature of cortical functional architecture linking microcircuit components with global brain networks.

## Introduction

The cerebral cortex orchestrates high-level brain functions ranging from perception and cognition to motor control, but the cellular basis of cortical network organization remains poorly understood. Across mammalian species, the cortex consists of dozens to over a hundred cortical areas, each featuring specific input-output connections with multiple other areas, thereby forming numerous functional subnetworks of information processing ^1,2^. Within each area, seminal discoveries have revealed columnar organization of neurons with similar functional properties ^3,4^, a foundational concept that has guided decades of cortical research ^5^. Across multiple cortical layers, diverse neuron types form intricate connections with each other and with neurons in other brain regions, constituting a “canonical circuit” that is duplicated and modified across areas ^6–8^. An enduring challenge is to decipher the cellular basis of cortical architecture characterized by such nested levels of organization that integrates microcircuits with global networks across spatiotemporal scales ^9,10^.

Among diverse cortical cell types, glutamatergic pyramidal neurons (PNs) constitute key elements for constructing the cortical architecture ^7,11^. Whereas PN dendrites and local axonal arbors form the skeleton of local cortical microcircuits, their long-range axons mediate communication with other cortical and subcortical regions. Indeed, PNs can be divided based on their long-range projections into hierarchically organized major classes and subclasses, each comprising finer grained projection types ^7,12^. One major class is the pyramidal tract (PT) neurons, which gives rise to corticofugal pathways that target all subcortical regions down to the brainstem and spinal cord. Another major class is the intratelencephalic (IT) neurons, which targets other cortical and striatal regions, including those in the contralateral hemisphere. Using this connectionist framework, recent studies in rodents have achieved a comprehensive description of cortical areal subnetworks ^13–15^ and have begun to reveal their cellular underpinnings ^16,17^. However, how these anatomically-defined networks relate to functional cortical networks remains unclear, as such studies require methods to monitor neural activity patterns in real time across large swaths of cortical territory yet with cell type resolution and in behaving animals.

Traditional fMRI methods measure brain-wide metabolic activities as a proxy of neural activity but with relatively poor spatial and temporal resolution ^18,19^. Conversely, single unit recording ^20^ and two-photon calcium imaging ^21^ achieve real-time monitoring of neural activity with cellular resolution, but typically with rather limited spatial coverage. Recent advances in widefield calcium imaging provide an opportunity to monitor neural activity in real time across a wide expanse of the mouse cortex at cellular resolution ^22^. However, most widefield studies to date have imaged activity either of broad neuronal classes ^23–33^ or of laminar subpopulations containing mixed projection types ^34–38^. Thus, while these studies have offered new insights into cortical activity during different brain states, they have yet to resolve and compare activity in distinct PN projection types. In particular, how IT and PT PNs each contribute to cortical processing during different brain states and as an animal moves and processes sensory information remains to be fully elucidated.

We have recently generated a comprehensive genetic toolkit for targeting the hierarchical organization of PNs in mouse cerebral cortex ^39^. This toolkit includes the *PlxnD1* and *Fezf2* driver lines that readily distinguish IT and PT PN types respectively. Here, we use these two driver lines to selectively express genetically-encoded calcium indicator in IT^PlxnD1^ and PT^Fezf2^ subpopulations and carried out widefield imaging across the dorsal cortex of head-fixed mice during a range of brain states. We found that IT^PlxnD1^ and PT^Fezf2^ show distinct activity dynamics during quiet wakefulness, spontaneous movements, upon sensory stimulation, and under anesthesia. Furthermore, we revealed distinct IT^PlxnD1^ and PT^Fezf2^ subnetworks dynamically tuned to different components of feeding behavior, including food retrieval, coordinated mouth-hand movements and food ingestion. Optogenetic inhibition of IT^PlxnD1^ and PT^Fezf2^ in key cortical areas in these subnetworks disrupted distinct components of this behavior. Anterograde tracing of IT^PlxnD1^ and PT^Fezf2^ from these key areas revealed projection patterns that contribute to functional activation patterns of corresponding subnetworks. Together, these results demonstrate that IT and PT subpopulations form parallel cortical processing streams and output pathways with spatiotemporal activity patterns that are distinct and change dynamically and differentially with behavioral state. Consequently, in addition to the concept of columnar organization, these experiments reveal that dynamic PN type subnetworks are a key feature of cortical functional architecture that integrates cortical microcircuits to global brain networks.

## Results

Although wide-field mesoscopic calcium imaging in rodents has the potential to reveal the cellular basis of functional cortical networks, realizing this potential critically depends on the resolution for distinguishing different PN types through specific expression of genetic encoded calcium sensors. To date, most if not all studies examined subpopulations (e.g. those defined by Thy1 and Rbp4 transgenic lines) each comprising mixtures of different projection classes ^35–38^, which precluded a direct and clear link between projection and function defined cortical networks. We have recently generated gene knockin mouse driver lines that enable specific genetic access to hierarchical PN classes and subpopulations within each class ^39^. Among these, the *Fezf2-CreER* line captures the large majority of PT population (PT^Fezf2^) in layer 5b and 6 (L5b/6), and the *PlxnD1-CreER* captures an IT subpopulation (IT^PlxnD1^) which resides in both layers 2/3 (L2/3) and layer 5a (L5a). Whereas PTs^Fezf2^ project to mostly ipsilateral subcortical regions and represent over 95% of corticospinal projection neurons, ITs^PlxnD1^ project to ipsi- and contra-lateral cortical and striatal targets ^39^. As IT is the largest and most diverse PN class comprising intracortical, callosal, and cortico-striatal PNs across layer 2-6, using double mRNA in situ hybridization we estimated that *PlxnD1* represent about 63% of IT neurons marked by *Satb2* (Fig. 1b). Quantification of cell distribution using serial two-photon tomography (STP) and CCF (mouse brain Common Coordinate Framework) registration demonstrated that PT^Fezf2^ and IT^PlxnD1^ are distributed across the entire dorsal neocortex (Fig, 1a, Extended data Fig. 13a)^39^. Therefore, *Fezf2-* and *PlxnD1*-driver lines represent specific and reliable tools that categorically distinguish the top-level PT and IT classes.

**Figure 1.**
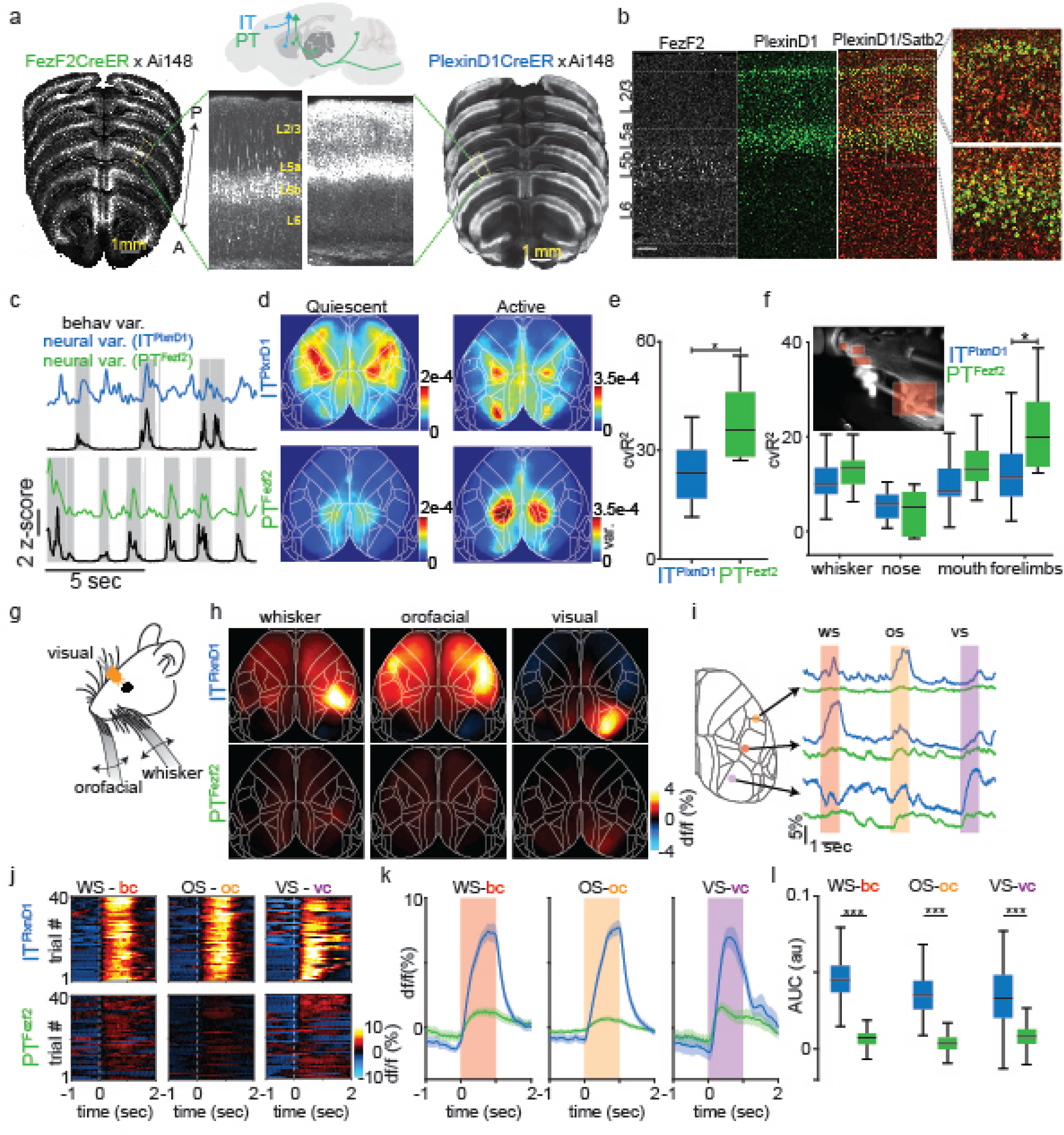
Distinct cortical activity patterns of IT^PlxnD1^ and PT^Fezf2^ during wakeful resting and upon sensory input. a. Serial two photon tomography images showing distribution of GCaMP6f labeled PT^Fezf2^ and IT^PlxnD1^ neurons across dorsal cortex in *FezF2-CreER;Ai148* and *PlexinD1-CreER:Ai148* mice respectively. Arrow indicates anterior (A) – posterior (P) axis. Yellow text in zoomed in image indicates approximate location of layer 2/3 (L2/3), layer 5a (L5a), layer 5b (L5b) and layer 6 (L6). Sagittal schematic depicts the major projection patterns of IT and PT neurons. b. mRNA in situ images and distribution of *Fezf2^+^* (left), *PlexinD1^+^* (middle) cells. Double in situ overlaid (right) shows *Satb2*^+^ in red and *PlexinD1*^+^ in green. Note that *PlexinD1*^+^ cells represent a subset of *Satb2^+^* IT cells. c. Example z-scored variance trace of behavior from video recordings (black trace) and corresponding z-scored variance trace of neural activity from IT^PlxnD1^ (blue) and PT^Fezf2^ (green) from wide-field GCaMP6f imaging. Gray blocks indicate active episodes identified from behavior variance trace. d. Average variance maps of spontaneous neural activity during active (right) and quiescent (left) episodes (12 sessions from 6 mice for each cell type). e. Distribution of the percentage of cross validated IT^PlxnD1^ and PT^Fezf2^ neural activity variance explained by full frame behavior variance obtained from a linear encoding model (12 sessions from 6 mice for each cell type, two-sided Wilcoxon rank sum test (*U=101, z=-2.8, p = 0.0051)).* f. Distribution of the percentage of cross validated IT^PlxnD1^ and PT^Fezf2^ neural activity variance explained by variance of specific body part movements obtained from the linear encoding model (12 sessions from 6 mice for each cell type, two-way ANOVA cell type*body part (*F_3,88_=3.21, p=0.03)* with Tukey-Kramer post hoc multiple comparison test IT^PlxnD1^ vs. PT^Fezf2^ whisker (*p=0.94),* IT^PlxnD1^ vs. PT^Fezf2^ nose (*p=0.99),* IT^PlxnD1^ vs. PT^Fezf2^ mouth (*p=0.7),* IT^PlxnD1^ vs. PT^Fezf2^ forelimbs (*p=0.007)).* g. Illustration of unimodal sensory stimulation paradigm. h. Mean activity maps of IT^PlxnD1^ and PT^Fezf2^ in response to corresponding unimodal sensory simulation (average of 12 sessions from 6 mice for each cell type). i. Single trial IT^PlxnD1^ and PT^Fezf2^ activity within orofacial (yellow), whisker (red) and visual (purple) areas during orofacial (os), whisker (ws) and visual (vs) stimulation. j. Single trial heat maps of IT^PlxnD1^ and PT^Fezf2^ activity from whisker (bc), orofacial (oc) and visual cortex (vc) in response to corresponding sensory stimulus from 1 example mouse for each cell type. k. Mean temporal dynamics of IT^PlxnD1^ and PT^Fezf2^ activity in whisker (bc), orofacial (oc) and visual cortex (vc) in response to corresponding sensory stimulus (average of 240 trials in 12 sessions from 6 mice for each cell type, shaded region indicates ±2 s.e.m). l. Distribution of IT^PlxnD1^ and PT^Fezf2^ activity intensity in whisker, orofacial and visual cortex in response to corresponding sensory stimulus (240 trials in 12 sessions from 6 mice for each cell type, two-way ANOVA cell type*stimulation (*F_2,1434_=38.28, p=0)* with Tukey-Kramer post hoc multiple comparison test IT^PlxnD1^ vs. PT^Fezf2^ whisker stimulation (*p=2e-8),* orofacial stimulation (*p=2e-8),* visual stimulation (*p=2e-8)*); *p<0.05, **p<0.005, ***p<0.0005.

### Distinct cortical activation patterns of IT^PlxnD1^ and PT^Fezf2^ during quiescent resting state and spontaneous movements

To examine IT^PlxnD1^ and PT^Fezf2^ activation patterns reflected by their calcium dynamics, we bred *PlxnD1* and *Fezf2* driver lines with a Cre-activated GCaMP6f reporter line (Ai148). Using serial two-photon tomography (STP), we revealed global GCaMP6f expression pattern for each cell type across the dorsal cortex. Consistent with previous results ^39^, GCaMP6f expression was restricted to L2/3 and 5a PNs in *PlxnD1* mice, and to L5b and to a lower extent in L6 PNs in *Fezf2* mice (Fig. 1a.). We first characterized network dynamics of IT^PlxnD1^ and PT^Fezf2^ during wakeful resting state in head-fixed mice using epifluorescence wide-field imaging to measure neuronal activity across the dorsal cortex (Extended data 1a, b). During this state, animals alternated between quiescence and spontaneous whisker, forelimb, and orofacial movements (Suppl. Videos 1, 2). We quantified full frame variance from behavioral videos recorded simultaneously with wide-field GCaMP6f imaging and identified episodes of quiescence and movements (active, Fig. 1c, Methods). We measured variance of neural activity for all pixels in each episode and found significantly higher variance during spontaneous movements versus quiescent episode in both cell types (Extended data 2b; quiescent vs. active variance median (1^st^ – 3^rd^ quartile), IT^PlxnD1^ 0.67e-4 (0.62e-4 – 0.72e-4) vs. 1.02e-4 (0.73e-4 – 1.31e-4), PT^Fezf2^ 0.33e-4 (0.28e-4 – 0.45e-4) vs. 0.59e-4 (0.49e-4 – 1.64e-4)). We then measured variance per pixel to obtain a cortical activation map during each episode. While both IT^PlxnD1^ and PT^Fezf2^ were active across broad areas, they differed significantly in strongly active areas (Fig. 1d, Extended data 2a,c,d). In quiescent episodes, IT^PlxnD1^ were most active across forelimb, hindlimb, and frontolateral regions while PT^Fezf2^ were only strongly active in posteromedial parietal areas. During spontaneous movement episodes, IT^PlxnD1^ showed strong activation in hindlimb and visual sensory areas while PT^Fezf2^ showed localized activation in the posteromedial parietal regions, with lower activation in other regions (Fig. 1d, Extended data 2a). We then compared the variance distribution at each pixel between the two populations (across mice and sessions) and used false discovery rate (FDR, q = 0.05) to identify pixels that were significantly different between the two conditions (Ext Data Fig. 2c). To measure consistency of resting state activity patterns we quantified Pearson’s correlation between spatial maps across mice and session and found strong correlation within and weak correlation between PN types during quiescent episodes (Extended data 2a,d; IT^PlxnD1 vs. PlxnD1^ vs. PT^Fezf2 vs. Fezf2^ vs. IT/PT^PlxnD1 vs. Fezf2^ correlation coefficient (r) median (1^st^ – 3^rd^ quartile), 0.65 (0.52 - 0.74) vs. 0.64 (0.53 – 0.71) vs. 0.22 (0.09 – 0.47)). During spontaneous movement (Active) episodes, IT^PlxnD1^ activity maps were far more variable compared to PT^Fezf2^ (Extended data 2a,e; IT^PlxnD1 vs. PlxnD1^ vs. PT^Fezf2 vs. Fezf2^ vs. IT/PT^PlxnD1 vs. Fezf2^ correlation coefficient (r) median (1^st^ – 3^rd^ quartile), 0.44 (0.31 – 0.58) vs. 0.81 (0.77 – 0.86) vs. 0.39 (0.18 – 0.55)). Principal component analysis on the combined spatial maps of both PN types revealed non-overlapping clusters across both episodes, indicating distinct cortical activation patterns between IT^PlxnD1^ and PT^Fezf2^ (Extended data 2f,g). To additionally visualize the magnitude of activation, we computed the 75^th^ and 95^th^ percentile of the activity at each pixel across states for both PN types and found them comparable to the variance maps (Extended data 2h)

To investigate the correlation between neural activity and spontaneous movements, we built a linear encoding model using the top 200 singular value decomposition (SVD) temporal components of the behavior video as independent variables to explain the top 200 SVD temporal components of cortical activity dynamics (Extended data 3a, b). The top 200 neural and behavior components explained more than 85 % of the variance in both IT^PlxnD1^ and PT^Fezf2^ with no significant difference in the amount of variance explained between the two populations (Extended data 3a). By measuring the proportion of 5-fold cross-validated activity variance explained by spontaneous movements (Methods), we found PT^Fezf2^ activity to be more strongly associated with spontaneous movements compared to IT^PlxnD1^ (Fig. 1e; IT^PlxnD1^ vs. PT^Fezf2^ cvR^2^ (%) median (1^st^–3^rd^ quartile), 23.7 (16.8 – 30.1) vs. 35.6 (28.2 – 46); see Methods). We then extracted mean time varying variance from parts of the behavior video frames covering nose, mouth, whisker, and forelimb regions (Fig. 1f inset) and identified the proportion of neural variance accounted by each body movement (Extended data 3c). While whisker and mouth contributed to some extent and equally between the two PN types, forelimbs contributed to significantly larger fraction of neural activity of PT^Fezf2^ than IT^PlxnD1^ (Fig. 1f, IT^PlxnD1^ vs. PT^Fezf2^ cvR^2^ (%) median (1^st^–3^rd^ quartile), whiskers 10 (7.9 – 13.4) vs. 13.4 (10 – 14.9), nose 5.8 (2.8 - 7.7) vs. 5.1 (−1.1 – 8.3), mouth 8.5 (7.5 – 13.3) vs. 13.2 (10.6 – 17.1), forelimbs 11.5 (7.5 – 16.4) vs. 19.9 (13.8 – 27.3); see Methods).

To evaluate whether or to what extent difference in PT^Fezf2^ and IT^PlxnD1^ responses could be explained by their difference in GCaMP6f expression we compared distribution of df/f values from all pixels during spontaneous behavior and found that both PNs show df/f values across a similar range (Extended data 4a). We also measured the peak df/f value at each pixel for every session during spontaneous behavior and compared their distribution (Extended data 4b). We found no regions (pixels) to be significantly different between the two populations (Extended data 4b,c). To verify if signal correction resulted in similar removal of artifacts, we performed df/f measurements with and without hemodynamic corrections. We then computed df/f measurements from the hindlimb sensory area and aligned them to the onset of spontaneous movements (Extended data 4d). We compared the difference in peak values (0 – 1 sec post movement onset) and Pearson’s correlation between corrected and uncorrected signals and found no significant difference between the two populations (Extended data 4e). Together, these results demonstrate distinct activation patterns of IT^PlxnD1^ and PT^Fezf2^ during wakeful resting state with preferential PT^Fezf2^ activation associated with spontaneous movements compared to IT^PlxnD1^.

### Unimodal sensory inputs preferentially activate IT^PlxnD1^ over PT^Fezf2^

We next investigated IT^PlxnD1^ and PT^Fezf2^ activation following sensory inputs of the somatosensory and visual system (Fig. 1g, Methods). We computed mean response per pixel across time for the stimulus duration (1 sec) associated with whisker, orofacial and visual stimulation to identify activated cortical areas (Fig. 1h.). Light stimulation and tactile stimulation of whisker and orofacial region strongly activated IT^PlxnD1^ in primary visual, whisker and mouth-nose somatosensory cortex, respectively, but resulted in no or weak activation of PT^Fezf2^ in those cortical areas (Fig 1h). To compare response intensities, we identified centers of maximum activation from the IT^PlxnD1^ spatial maps and extracted temporal signals within a circular window around the centers and computed single trial dynamics and trial averaged responses from both PN types (Fig. 1i-k). Again, we found strong IT^PlxnD1^ activation but weak PT^Fezf2^ activation in primary sensory cortices in response to whisker, orofacial, and visual stimulus (Fig. 1j,k). We then measured response intensities by integrating the signal for the stimulus duration and found significantly higher activity in IT^PlxnD1^ compared to PT^Fezf2^ (Fig. 1l; IT^PlxnD1^ vs. PT^Fezf2^ AUC (au) median (1^st^ – 3^rd^ quartile), whisker stimulation 0.045 (0.037-0.054) vs. 0.007 (0.003 – 0.01), orofacial stimulation 0.035 (0.026 – 0.044) vs. 0.003 (0 – 0.008), visual stimulation 0.033 (0.02 – 0.077) vs. 0.008 (0.003 – 0.013)). To visualize sensory responses across cortical regions irrespective of response intensity, we peak normalized the maps and found activation in corresponding sensory cortices in PT^Fezf2^ along with broader activation including the retrosplenial areas (Extended data 4f). To measure consistency in activation patterns we computed Pearson’s correlation between the spatial maps across mice and sessions and found stronger correlations between IT^PlxnD1^ spatial maps compared to PT^Fezf2^ (Extended data 4g; IT^PlxnD1 vs. PlxnD1^ vs. PT^Fezf2 vs. Fezf2^ vs. IT/PT^PlxnD1 vs. Fezf2^ correlation coefficient (r) median (1^st^ – 3^rd^ quartile), whisker stimulation 0.77 (0.73 – 0.82) vs. 0.49 (0.26 – 0.59) vs. 0.37 (0.22 – 0.54), orofacial stimulation 0.86 (0.8 – 0.89) vs. 0.31 (0.05 – 0.57) vs. 0.50 (0.24 – 0.99), visual stimulation 0.84 (0.8 – 0.87) vs. 0.56 (0.40 – 0.71) vs. 0.63 (0.55 – 0.71)). Considering that ITs^PlxnD1^ constitute a major subpopulation of ITs (Figure 1b), these results provide the first set of in vivo evidence that sensory inputs predominantly activate IT compared to PT PNs, consistent with previous findings that thalamic input predominantly impinges on IT but not PT cells ^40^. Notably, ITs^PlxnD^ activation per se does not lead to significant PT activation at the population level despite demonstrated synaptic connectivity from IT to PT PNs in cortical slice preparations ^41^. It is possible that ITs^PlxnD^ to PT^Fezf2^ synaptic efficacy is modulated by brain states or that another IT subpopulation might more directly activate PT^Fezf2^.

To confirm widefield responses reflected calcium dynamics at cellular resolution, we used two photon imaging to measure neural responses from single IT^PlxnD1^ and PT^Fezf2^ neurons from contralateral barrel cortex in response to whisker stimulation (Extended data 5a, Methods). We recorded from cell bodies of IT^PlxnD1^ and apical dendrites of PT^Fezf2^ (dendritic calcium activity in layer 5B is strongly correlated to cell body dynamics^42–46^, Extended data 5b) and used linear modelling approach to classify neurons as activated, inhibited or unclassified groups (Methods). We first measured the average response for each group of neurons form a single field of view (FOV) and found that [within the activated group] IT^PlxnD1^ neurons showed significantly higher response compared to PT^Fezf2^ with very little difference among the inhibited group during whisker stimulation (Extended data 5c,d). Average response of all neurons combined from a single FOV resulted in IT^PlxnD1^ displaying significantly larger response compared to PT^Fezf2^ (Extended data 5e). Combining neuronal responses from all mice across all FOV’s resulted in similar response characteristics (Extended data 5 f-h). These responses followed dynamics very similar to those observed using widefield imaging during whisker stimulation (Fig. 1k). Additionally, a larger proportion of IT^PlxnD1^ neurons were activated in response to whisker stimulation compared to PT^Fezf2^ (across FOVs IT^PlxnD1^ vs. PT^Fezf2^ mean ± s.d. (%), activated: 36.8 ± 15.5 vs. 28.1 ± 12.7, inhibited: 50.8 ± 20.6 vs. 44.8 ± 11.7, unclassified: 44.1 ± 12.3 vs. 54.9 ±15.6; data from all cells combined in Extended data 5i). These results show that the response properties observed in widefield imaging of IT^PlxnD1^ and PT^Fezf2^ neurons closely reflect their cellular resolution dynamics.

### Distinct PT^Fezf2^ and IT^PlxnD1^ subnetworks tuned to different sensorimotor components of a feeding behavior

To examine the activation patterns of IT^PlxnD1^ and PT^Fezf2^ during sensorimotor processing, we designed a head-fixed setup that captures several features of natural mouse feeding behavior. In this setup, mice sense a food pellet approaching from the right side on a moving belt, retrieve pellet into mouth by licking, recruit both hands to hold the pellet and initiate repeated bouts of hand-mouth coordinated eating movements that include: bite while handling the pellet, transfer pellet to hands while chewing, raise hands to bring pellet to mouth thereby starting the next bout (Fig. 2a, Suppl video 3). We used DeepLabCut ^47^ to track pellet and body parts in video recordings and wrote custom algorithms to identify and annotate major events in successive phases of this behavior (Methods, Fig. 2b, Extended data 6a, Suppl. Video 3).

**Figure 2.**
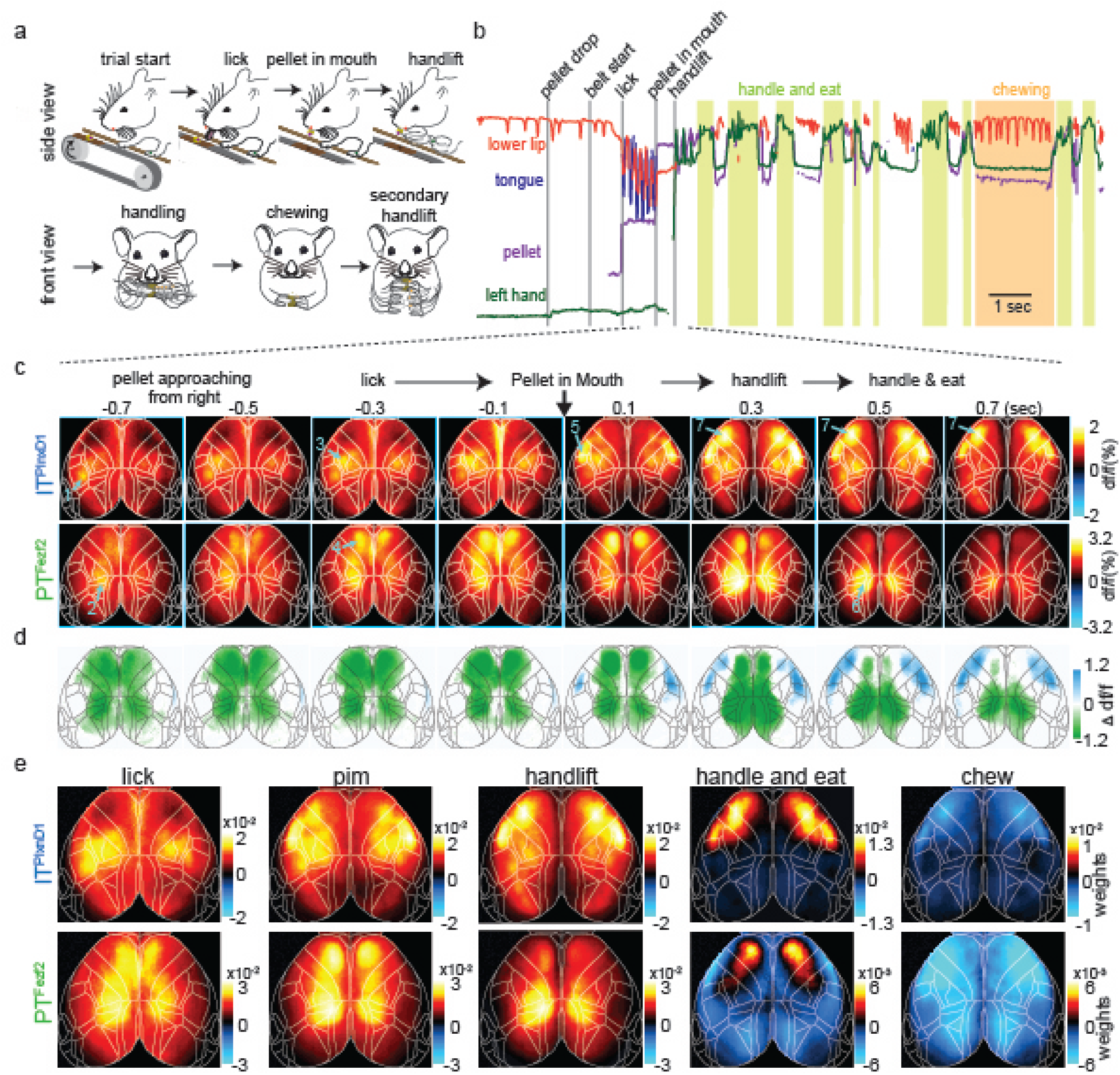
Distinct PT^Fezf2^ and IT^PlxnD1^ subnetworks tuned to different sensorimotor components of a feeding behavior. a. Schematic of the head-fixed feeding behavior showing the sequential sensorimotor components. b. Example traces of tracked body parts and episodes of classified behavior events. Colored lines represent different body parts as indicated (light green shade: handle-and-eat episodes; orange shade: chewing episode). c. Mean IT^PlxnD1^ and PT^Fezf2^ sequential activity maps during the feeding sequence before and after pellet-in-mouth (PIM) onset (IT^PlxnD1^ – 23 sessions from 6 mice, PT^Fezf2^ – 24 sessions from 5 mice). Note the largely sequential activation of areas and cell types indicated by arrows and numbers: 1) left barrel cortex (IT^PlxnD1^) when right whiskers sensed approaching pellet; 2) parietal node (PT^Fezf2^) while making postural adjustments as pellet arrives; 3) forelimb sensory area (IT^PlxnD1^) with limb movements that adjusted grips of support bar as pellet approaches closer; 4) frontal node (PT^Fezf2^) during lick; 5) orofacial sensory areas (FLP (Frontolateral Posterior), IT^PlxnD1^) when pellet-in-mouth; 6) parietal node again during hand lift; 7) FLA (Frontolateral Anterior)-FLP (IT^PlxnD1^) on handling and eating the pellet. d. Difference between IT^PlxnD1^ and PT^Fezf2^ average activity maps at each time step as in panel c. Only significantly different pixels are displayed (*two-sided Wilcoxon rank sum test with p-value adjusted by FDR = 0.05).* Blue pixels indicate values significantly larger in IT^PlxnD1^ compared to PT^Fezf2^ and vice versa for green pixels. e. Spatial maps of IT^PlxnD1^ (top) and PT^Fezf2^ (bottom) regression weights from an encoding model associated with lick, PIM, hand lift, eating and handling, and chewing (IT^PlxnD1^ - 23 sessions from 6 mice, PT^Fezf2^ - 24 sessions from 5 mice).

We then imaged the spatiotemporal activation patterns of IT^PlxnD1^ and PT^Fezf2^ across dorsal cortex while mice engage in the behavior (Suppl. Videos 4, 5). We calculated average GCaMP6f signals per pixel in frames taken at multiple time points centered around when mice retrieve pellet into the mouth (pelletin-mouth, or PIM) (Fig. 2c, frames displayed at 200ms time steps). Upon sensing the pellet approaching from the right side, mice adjust their postures and hand grip of the support bar while initiating multiple right-directed licks until retrieving the pellet into the mouth. During this period (Fig.2c, −0.7-0 seconds), IT^PlxnD1^ was first activated in left whisker primary sensory cortex (SSbfd), which then spread to bi-lateral forelimb and hindlimb sensory areas (Fig 2c, Suppl. Video 4). Almost simultaneously or immediately after, PT^Fezf2^ was strongly activated in left medial parietal cortex (parietal node) just prior to right-directed licks, followed by bi-lateral activation in frontal cortex (medial secondary motor cortex, frontal node) during lick and pellet retrieval into mouth (Fig 2c, Suppl. Video 5). During the brief PIM period when mice again adjusted postures and then lift both hands to hold the pellet (Fig 2c, 0-0.1 sec), IT^PlxnD1^ was activated in bi-lateral orofacial primary sensory cortex (frontolateral posterior node; FLP) and subsequently in an anterior region spanning the lateral primary and secondary motor cortex (frontolateral anterior node; FLA), while PT^Fezf2^ activation shifted from bilateral frontal to parietal node. In particular, PT^Fezf2^ activation in parietal cortex reliably preceded hand lift. Following initiation of repeated bouts of eating actions, IT^PlxnD1^ was prominently activated in bilateral FLA and FLP specifically during coordinated oral-manual movements such as biting and handling, whereas PT^Fezf2^ activation remained minimum throughout the dorsal cortex. For each time point, we compared the average df/f values at each pixel to identify regions that were significantly different between the two populations, which further confirmed the differential flow of activation pattern described earlier (Fig. 2d). To examine the reliability of the activation patterns, we measured Pearson’s correlation within and between activation maps of the two populations across mice and sessions at each time point (Extended data 6b). While PT^Fezf2^ activity maps strongly correlated with lick and handlift, IT^PlxnD1^ activity maps showed a sharp increase following pellet in mouth which continued during oro-manual handling. We found rather weak correlation between activation patterns of the two populations across time (Extended data 6b).

To obtain a spatial map of cortical activation during specific feeding events, we build a linear encoding model using binary time stamps associated with the duration of lick, PIM, hand lift, handling, and chewing as independent variables to explain the top 200 SVD temporal components of neural activity. We then transformed the regression weights obtained from the model to the spatial domain of neural activity using SVD spatial components to obtain a cortical map of weights associated with each event (Fig. 2e, Methods). This analysis largely confirmed and substantiated our initial observations (Fig. 2c). Indeed, during lick onset IT^PlxnD1^ was active in left barrel cortex and bilateral forelimb/hindlimb sensory areas, which correlated with sensing the approaching pellet with contralateral whiskers and limb adjustments, respectively. In sharp contrast, PT^Fezf2^ was active along a medial parietal-frontal axis, which correlated with licking. During PIM, IT^PlxnD1^ was preferentially active in FLP with lower activity in FLA as well as in forelimb/hindlimb sensory areas, while PT^Fezf2^ was still active along the parietofrontal regions. Prior to and during hand lift, PT^Fezf2^ showed strong activation specifically in bilateral parietal areas, whereas IT^PlxnD1^ showed predominant bilateral activation in both FLA and FLP. During eating and pellet handling, IT^PlxnD1^ was strongly active in FLA and less active in FLP, while PT^Fezf2^ was only weakly active specifically in the frontal node (Fig 2e, note scale change in panels). Both IT^PlxnD1^ and PT^Fezf2^ showed significantly reduced activity across dorsal cortex during chewing (Fig. 2e).

To capture the most prominently activated cortical areas associated with onset of various feeding movements, we computed average activity per pixel during the progression from licking, retrieving pellet into mouth, to hand lift across mice and sessions (from 1 second before to 2 seconds after PIM, Fig 3a,d). We then compared the average df/f distribution at each pixel to identify regions that were significantly different between the two populations (Extended data 7a). This analysis confirmed that, while both cell types were activated across multiple regions, PT^Fezf2^ activation was most prominent along a medial parietal-frontal network whereas IT^PlxnD1^ was most strongly engaged along a frontolateral FLA-FLP network (Fig. 3a, d, Extended data 7a). To examine the reliability of the activation patterns, we computed Pearson’s correlation within and between the average activation maps of the two populations across mice and sessions. Activation patterns were much more strongly correlated within each population than between the two populations, with PT^Fezf2^ maps being more consistent than IT^PlxnD1^ (Extended data 7b). Principal component analysis on the combined spatial maps of both PN types revealed non-overlapping clusters indicating distinct cortical activation patterns between IT^PlxnD1^ and PT^Fezf2^ during various feeding movements (Extended data 7c).

**Figure 3.**
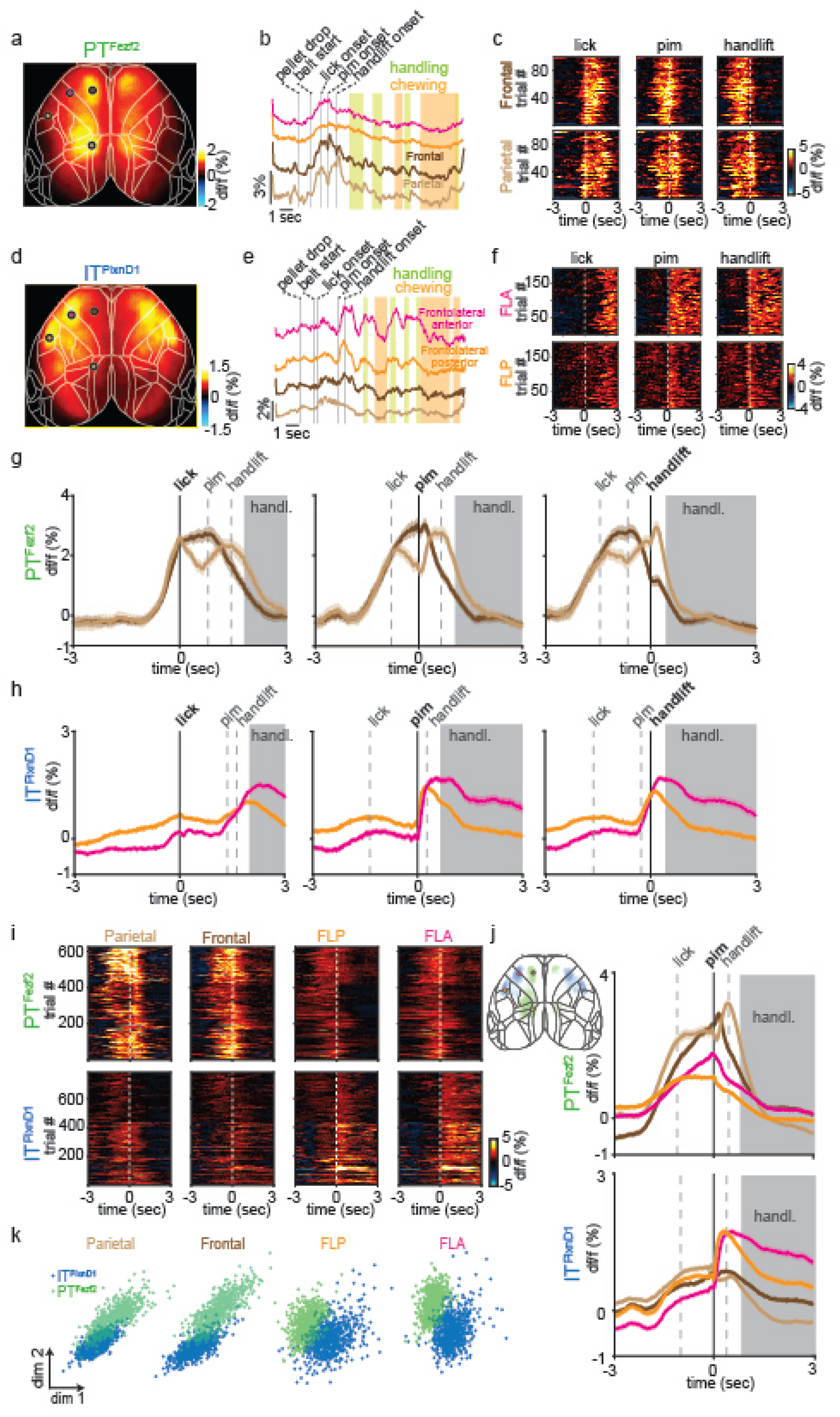
IT^PlxnD1^ and PT^Fezf2^ within frontolateral and parietofrontal nodes show distinct temporal dynamics during feeding behavior. a. Mean activity map of PT^Fezf2^ across the dorsal cortex during the feeding sequence from 1 second before to 2 seconds after PIM onset (24 sessions from 5 mice). b. Example single PT^Fezf2^ activity traces from FLA (magenta), FLP (orange), frontal (dark brown) and parietal (light brown) nodes during feeding behavior; vertical bars indicate different behavior events. c. Single trial heat maps of PT^Fezf2^ activities from frontal and parietal nodes centered to onset of licking, PIM, and hand lift (5 sessions from one example mouse). d. Mean activity map of IT^PlxnD1^ across the dorsal cortex during the feeding sequence from 1 second before to 2 seconds after PIM onset (23 sessions from 6 mice). e. Example single IT^PlxnD1^ activity traces from FLA, FLP, frontal and parietal nodes during feeding behavior. f. Single trial heat maps of IT^PlxnD1^ activities from FLA and FLP nodes centered to onset of licking, PIM and hand lift (5 sessions from one example mouse). g. Mean PT^Fezf2^ activity within frontal and parietal node centered to onset of lick, PIM, and hand lift. Grey dashed lines indicate median onset times of other events relative to centered event (5 sessions from one example mouse, shading around trace ±2 s.e.m). Grey shade indicates eating-handling episode. h. Mean IT^PlxnD1^ activity within FLA and FLP centered to onset of lick, PIM, and hand lift; grey dashed lines indicate median onset times of other events relative to centered event (5 sessions from one example mouse, shading around trace ±2 s.e.m). Grey shade indicates eating-handling episode. i. Single trial heat maps of IT^PlxnD1^ and PT^Fezf2^ activities within parietal, frontal, FLP and FLA nodes centered to PIM (IT^PlxnD1^ - 23 sessions from 6 mice, PT^Fezf2^ - 24 sessions from 5 mice). j. Mean IT^PlxnD1^ and PT^Fezf2^ temporal dynamics within parietal (light brown), frontal (dark brown), FLP (orange) and FLA (magenta) centered to PIM. Grey dashed lines indicate median onset times of lick and hand lift relative to PIM (IT^PlxnD1^ - 23 sessions from 6 mice, PT^Fezf2^ - 24 sessions from 5 mice, shading around trace ±2 s.e.m). Left inset: Overlaid activity maps of IT^PlxnD1^ (blue) and PT^Fezf2^ (green) after thresholding indicating distinct nodes preferentially active during the feeding sequence. k. Distribution of IT^PlxnD1^ and PT^Fezf2^ activities centered at PIM onset from parietal, frontal, FLP and FLA node and projected to the subspace spanned by the first two linear discriminant analysis dimensions (IT^PlxnD1^ - 23 sessions from 6 mice, PT^Fezf2^ - 24 sessions from 5 mice).

To characterize the temporal activation patterns of these key cortical nodes during behavior, we extracted temporal traces from the center within each of these 4 areas (averaged across pixels within a 280 μm radius circular window) and examined their temporal dynamics aligned to the onset of lick, PIM, and hand lift (Fig 3b, c, e, f). PT^Fezf2^ activity in the frontal node rose sharply prior to lick, sustained for the duration of licking until PIM, then declined prior to hand lift; on the other hand, PT^Fezf2^ activity in the parietal node increased prior to lick then declined immediately after, followed by another sharp increase prior to hand lift then declined again right after. In contrast, IT^PlxnD1^ activities in FLA and FLP did not modulate significantly during either licking or hand lift but increased specifically only when mice first retrieved pellet into mouth; while activation in FLP decreased following a sharp rise after pellet-in-mouth, activities in FLA sustained during biting and handling (Fig. 3g, h). To examine cortical dynamics from both populations within the same region, we measured GCaMP6f signals centered to PIM from all four nodes for each cell type. PT^Fezf2^ showed strong activation within parietal and frontal nodes specifically during lick and hand lift, whereas IT^PlxnD1^ showed significantly lower activation within these nodes during these episodes (Fig. 3i, j). In sharp contrast, IT^PlxnD1^ was preferentially active in FLA and FLP specifically during PIM with sustained activity especially in FLA during biting and handling, but no associated activity was observed in PT^Fezf2^ within these nodes during the same period (Fig. 3i,j). Similar difference in dynamics was observed from activity aligned to either lick or handlift onset (Extended data 7d)

To validate the differential temporal dynamics between PN types we performed linear discriminant analysis (LDA) on combined single trials of IT^PlxnD1^ and PT^Fezf2^ in each node (Methods). We trained an LDA model to discriminate if single trial activity from a node was associated with either IT^PlxnD1^ or PT^Fezf2^ and projected traces onto the top two dimensions identified by LDA to visualize the spatial distribution of projected clusters. This analysis revealed that IT^PlxnD1^ and PT^Fezf2^ activity clustered independently with little overlap, indicating that these PNs follow distinct temporal dynamics within each region (Fig. 3k). Altogether, these results indicate that IT^PlxnD1^ and PT^Fezf2^ operate in distinct and partially parallel cortical subnetworks, which are differentially engaged during specific sensorimotor components of an ethologically relevant behavior. It is important to note that the IT class comprises diverse subpopulations beyond IT^PlxnD1^; it is possible that activity of another IT subpopulation might more closely correlate with PT neurons.

### Feeding without hand occludes parietal PT^Fezf2^ activity

To investigate if the observed PN neural dynamics were causally related to features of the feeding behavior, we developed a variant of the feeding task in which mice lick to retrieve food pellet but eat without using hands (Fig. 4a); this was achieved by using a blocking plate to prevent hand lift until mice no longer attempted to use their hands during eating. We then measured IT^PlxnD1^activity in FLA-FLP nodes and PT^Fezf2^ activity in parietal-frontal nodes aligned to the onset of lick and PIM, and compared activity dynamics to those during normal trials in the same mice when they used hands to eat (Fig. 4b-e, Extended data 8a,b). Whereas PT^Fezf2^ activity in the frontal node did not show a difference with or without hand lift (Fig 4c, Extended data 8a, b), PT^Fezf2^ activity in parietal node showed a significant decrease in trials without hand lift specifically during the time when mice would have lifted hand during normal feeding trials (Fig 4c, Extended data 8a, b). On the other hand, IT^PlxnD1^ activity in FLA and FLP did not change for the duration from retrieving pellet through hand lift in trials with no hand lift compared with those in normal trials with hand lift. However, IT^PlxnD1^ activities in FLA and FLP showed a notable reduction during the eating-handling phase (Fig. 4d, e, Extended data 8a, b). We then computed integral of activity from PIM onset to 1 second after and compared results between trials with and without hand lift. Only PT^Fezf2^ activities in parietal node showed a sharp decline whereas IT^PlxnD1^ activities across FLA and FLP did not change significantly during the hand lift phase (Fig. 4f,g; PT^Fezf2^ Parietal lift vs. no lift activity intensity (au) median (1^st^ – 3^rd^ quartile), 0.023 (0.017 – 0.03) vs. 0.014 (0.008 – 0.019), PT^Fezf2^ Frontal lift vs. no lift, 0.018 (0.013 – 0.023) vs. 0.018 (0.014 – 0.022), IT^PlxnD1^ FLP lift vs. no lift 0.013 (0.01 – 0.017) vs. 0.013 (0.009 – 0.019), IT^PlxnD1^ FLA lift vs. no lift 0.014 (0.01 – 0.018) vs. 0.015 (0.012 – 0.019)).

**Figure 4.**
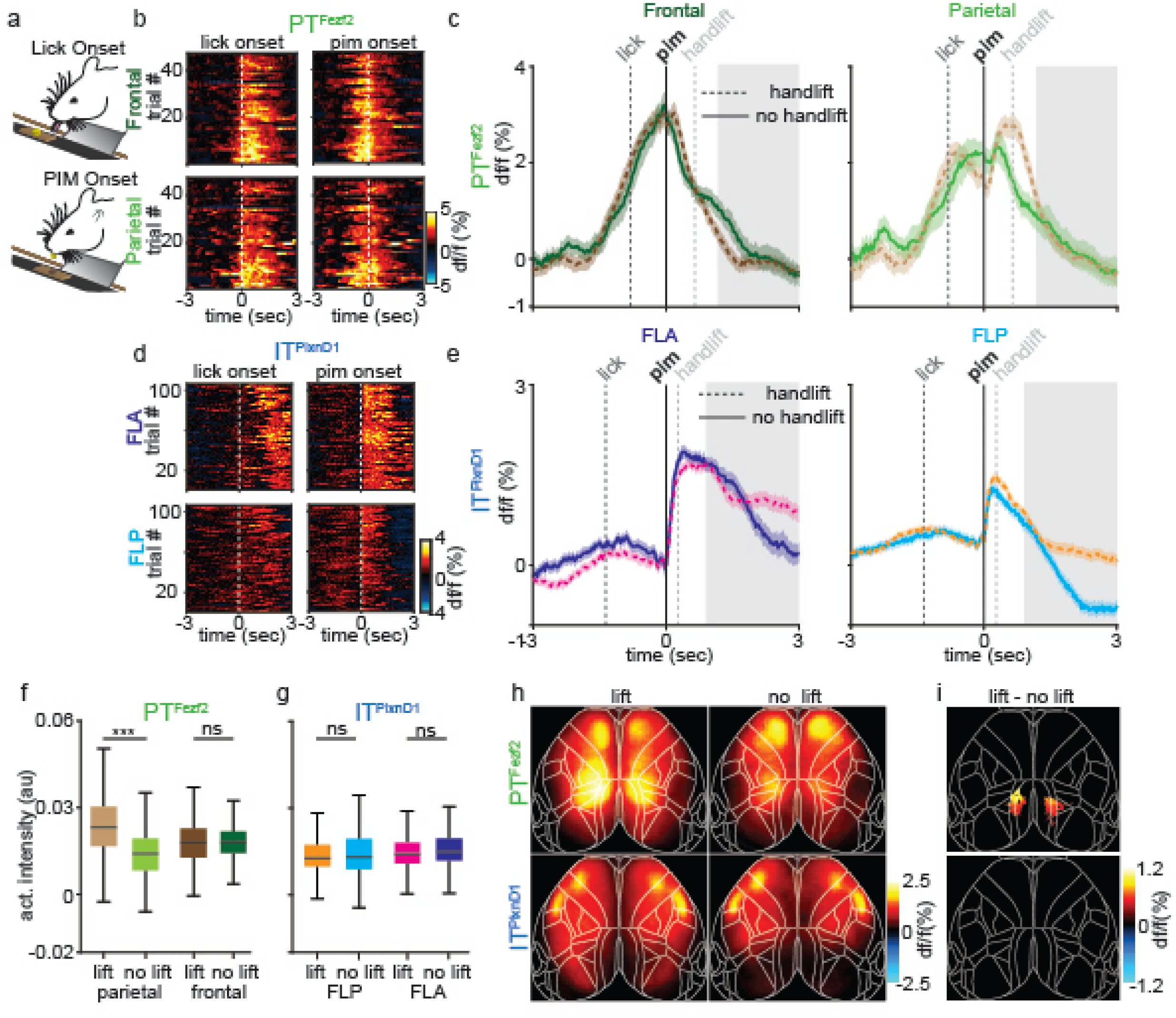
Feeding without hand lift selectively occludes PT^Fezf2^ activity in parietal node. a. Schematic of feeding without hand following preventing hand lift with a blocking plate. b. Single trial heat maps of PT^Fezf2^ activity within frontal and parietal nodes centered to onset of licking and PIM during feeding without hand lift (2 sessions from one example mouse). c. Mean PT^Fezf2^ activity within frontal (left) and parietal (right) nodes centered to onset of PIM during feeding with (frontal: dark brown, parietal: light brown, 5 sessions from one example mouse, shading around trace ±2 s.e.m) and without handlift (frontal: dark green, parietal: light green, 2 sessions from the same example mouse, shading around trace ±2 s.e.m). Grey shade indicates eating-handling episode during handlift sessions. d. Single trial heat maps of IT^PlxnD1^ activity within FLA and FLP centered to onset of licking and PIM during feeding without hand lift (3 sessions from one example mouse). e. Mean IT^PlxnD1^ activity within frontolateral anterior (left) and frontolateral posterior (right) nodes centered to onset of PIM during feeding with (FLA: magenta, FLP: orange, 5 sessions from one example mouse, shading around trace ±2 s.e.m) and without hand lift (FLA: dark blue, FLP: cyan, 3 sessions from the same example mouse, shading around trace ±2 s.e.m). Grey shade indicates eating-handling episode during handlift sessions. f. Distribution of PT^Fezf2^ activity intensity during 1 sec post PIM onset from parietal and frontal nodes across behaviors with and without hand lift (no lift: 440 trials in 13 sessions from 5 mice, lift: 647 trials in 24 sessions from the same 5 mice, two-way ANOVA region*lift *(F_1,2170_=158.93, p=0)* with Tukey-Kramer post hoc multiple comparison test parietal lift vs. no lift (*p=3.4e-8),* frontal lift vs. no lift (*p=0.99)).* g. Distribution of IT^PlxnD1^ activity intensity during 1 sec post PIM onset from FLA and FLP across behaviors with and without hand lift (no lift: 544 trials in 15 sessions from 6 mice, lift: 781 trials in 23 sessions from the same 6 mice, two-way ANOVA region*lift *(F_1,2646_=0.51, p=0.47)* with Tukey-Kramer post hoc multiple comparison test FLP lift vs. no lift (*p=0.43),* FLA lift vs. no lift (*p=0.06)).* h. Mean PT^Fezf2^ (top) and IT^PlxnD1^ (bottom) spatial activity maps during 1 sec post PIM onset for behaviors with (left) and without (right) hand lift. i. Difference in PT^Fezf2^ (top) and IT^PlxnD1^ (bottom) mean spatial activity maps between eating with and without hand lift. Only significantly different pixels are displayed (*twosided Wilcoxon rank sum test with p-value adjusted by FDR = 0.05).* Note that parietal areas in PT^Fezf2^ and no pixels in IT^PlxnD1^ are significantly different between lift and no lift.***p<0.0005.

To visualize cortical regions differentially modulated between feeding trials with or without hand-lift, we computed mean pixel-wise activity during a 1 second period after PIM onset from both trial types and subtracted the spatial map of no-hand-lift trials from that of hand-lift trials (Fig. 4h). We compared the df/f distribution at each pixel between the lift and no lift average maps across mice and sessions and only displayed significantly different pixels across the mean difference map (Fig. 4i). As expected only the parietal region in PT^Fezf2^ showed significantly higher activity during hand-lift compared to no-hand-lift trials while no pixels were significantly different within IT^PlxnD1^ (Fig. 4h, i). These results strengthen the correlation between parietal PT^Fezf2^ activation and hand lift movement during feeding; they also suggest that IT^PlxnD1^ activity in FLA and FLP is in part related to orofacial sensorimotor components of feeding actions.

### Inhibition of parietofrontal and frontolateral regions differentially disrupts sensorimotor components of feeding behavior

To investigate if the cortical regions active during feeding behavior was necessary for its proper execution, we adopted a laser scanning method to optogenetically inhibit bilateral regions of dorsal cortex using vGat-ChR2 mice expressing Channelrhodopsin-2 in GABAergic inhibitory neurons (Extended data. 9a). We examined the effects of bilateral inhibition of parietal, frontal, FLP and FLA nodes on different components of the behavior including pellet retrieval by licking, hand lift after PIM, and mouth-hand mediated eating bouts. We pooled the data from FLA and FLP as we did not observe major differences in disrupting each of the two areas.

During the pellet retrieval phase, inhibition of frontal and the frontolateral nodes during licking resulted in a sharp decrease of tongue extension, which recovered on average after about 0.5 seconds (Extended data 9b, c, before vs. during inhibition tongue length (au) median (1^st^ – 3^rd^ quartile), frontal 6.52 (0 – 10.08) vs. 4.54 (1.1 – 6.85), parietal 8.95 (1.47 – 13.37) vs. 7.65 (4.25 – 10.79), frontolateral 3.89 (0 – 9.15) vs. 1.4 (0 – 3.6)); inhibition also led to a significant delay and disruption in pellet retrieval to mouth (Extended data 9d; control vs. inhibition pim onset time (sec) median (1^st^ – 3^rd^ quartile), frontal 4.81 (4.66 – 5.12) vs. 5.07 (4.82 – 5.85), parietal 4.79 (4.66 – 5.13) vs. 4.93 (4.66 – 5.26), frontolateral 4.77 (4.56 – 5.12) vs. 5.07 (4.61 – 10.96)). These effects were not observed when inhibiting the parietal node (Extended data 9b-d). During the hand-lift phase after PIM, inhibition of frontal and frontolateral nodes prior to hand lift onset led to substantial deficit in the ability to lift hands towards mouth, resulting in a sharp decrease in the number of hand lift (Extended data 9e); notably, in trials when animals did initiate hand lift despite inhibition, hand trajectories appeared smooth and normal. Upon inhibiting the parietal node, on the other hand, no significant decrease of hand lift numbers per se was observed but there were substantial deficits in hand lift trajectory, characterized by erratic and jerky movements (Extended data 9f). To quantify these deficits, we measured the absolute value of the first derivative of 15 Hz low pass filtered (to remove high frequency noise) hand lift trajectory to obtain the time varying velocity during the lift episode and compared its integral between 0 to 1 second after hand lift between inhibition and control trials. While frontal and frontolateral inhibition did not affect lift trajectories among the successful trials, parietal inhibition resulted in a significant modulation of velocity during lift, indicating deficits in forelimb movements (Extended data 9f,g; control vs. inhibition area under curve (au) median (1^st^ – 3^rd^ quartile), frontal 1.36 (1.02 – 1.69) vs. 1.29 (0.87 – 1.68), parietal 1.34 (1.11 – 1.79) vs. 1.78 (1.25 – 2.35), frontolateral 1.39 (1.1 – 1.59) vs. 1.43 (1.12 – 1.77)).

During eating bouts with coordinated mouth-hand movements, after the initial hand lift from support bar to hold pellet in mouth, inhibiting the frontal and frontolateral nodes severely impeded mice’s ability to bring pellet to mouth (i.e. secondary hand lifts when mice bring pellet to mouth after holding it while chewing); this deficit recovered immediately after the release of inhibition. Inhibiting the parietal node resulted in only a slight disruption of these secondary hand lifts (Extended data 9h). To quantify these deficits, we measured the average distance of left fingers to the mouth and the duration of the hand held close to mouth. We found significant deficits in all these parameters in inhibition trials for all cortical areas compared to those of control trials (Extended data 9i-j; control vs. inhibition finger-mouth distance (au) median (1^st^ – 3^rd^ quartile), frontal 0.15 (0.13 – 0.17) vs 0.18 (0.16 – 0.21), parietal 0.14 (0.11 – 0.18) vs. 0.16 (0.13 – 0.19), frontolateral 0.14 (0.11 – 0.17) vs. 0.2 (0.17 – 0.23), control vs. inhibition hand at mouth duration (sec) median (1^st^ – 3^rd^ quartile), frontal 1.85 (1.44 – 2.27) vs. 1.07 (0.44 – 1.27), parietal 1.74 (1.27 – 2.2) vs 1.48 (0.93 – 1.93), frontolateral 2.25 (1.76 – 2.59) vs. 1.22 (0.83 – 1.66)). These results show that the parietofrontal and frontolateral regions are necessary to orchestrate orofacial and forelimb movements that enable pellet retrieval and mouth-hand coordinated eating behavior.

### Inhibiting IT^PlxnD1^ and PT^Fezf2^ in frontal and frontolateral regions disrupts distinct components of feeding behavior

We then investigated if IT^PlxnD1^ and PT^Fezf2^ neurons within the same cortical region were causally associated with distinct sensorimotor components of the feeding behavior. We expressed-light activated Anion Channelrhodopsin (GtACR1) specifically within IT^PlxnD1^ or PT^Fezf2^ neurons in frontal and frontolateral regions of the same mouse and examined the effects of bilaterally inhibiting either population during specific phases of the feeding behavior (Methods, Fig. 5a). During the pellet retrieval phase, inhibition of PT^Fezf2^ neuron in both frontal and frontolateral nodes resulted in a sharp decrease in tongue length throughout the inhibition whereas disrupting IT^PlxnD1^ neurons resulted in a momentary decrease at inhibition onset after which the animal recovered immediately to lick the pellet (Fig. 5b, Suppl. Video 6,7). This resulted in a significant decrease in the mean tongue length during inhibition compared to control trials only when disrupting PT^Fezf2^ but not IT^PlxnD1^ neurons (Fig 5c, mean tongue length (au) control vs. inhibition median (1^st^ – 3^rd^ quartile), IT^PlxnD1^ frontal 0.058 (0.017 – 0.09) vs. 0.054 (0.027 – 0.088), IT^PlxnD1^ FLA 0.044 (0.009 – 0.08) vs. 0.055 (0.029 – 0.093), PT^Fezf2^ frontal 0.044 (0.006 – 0.092) vs. 0.023 (0.003 – 0.059), PT^Fezf2^ FLA 0.025(0.003 – 0.084) vs. 0.01 (0.003 – 0.038)). Inhibiting PT^Fezf2^ activity in both frontal and frontolateral regions, after the mouse picked the pellet, strongly affected the ability to bring hands close to the mouth to hold the pellet, resulting in a sharp decrease in the proportion of handlift episodes while only a small effect was observed on inhibiting IT^PlxnD1^ neurons (Fig. 5d, Suppl. Video 8). Disrupting PT^Fezf2^ activity after retrieving the pellet and bringing hands to mouth (during food handling) strongly affected the gross mobility of hands such that mice were unable to properly bring the pellet towards the mouth (Fig 5e, Suppl. Video 9). To quantify this, we measured the average distance of the first finger to the mouth and its absolute velocity across time and found a significant increase in the hand-mouth distance and decrease in velocity during inhibition of PT^Fezf2^ neurons in both frontal and frontolateral regions (Fig. 5e,f,g hand-mouth distance (au) control vs. inhibition median (1^st^ – 3^rd^ quartile), IT^PlxnD1^ frontal 0.17 (0.14 – 0.19) vs. 0.16 (0.13 – 0.19), IT^PlxnD1^ FLA 0.17 (0.15 – 0.2) vs. 0.17 (0.13 – 0.2), PT^Fezf2^ frontal 0.14 (0.11 – 0.18) vs. 0.16 (0.1 – 0.28), PT^Fezf2^ FLA 0.13 (0.11 – 0.18) vs. 0.23 (0.14 – 0.34); hand velocity (au) control vs. inhibition median (1^st^ – 3^rd^ quartile), IT^PlxnD1^ frontal 0.006 (0.005 – 0.008) vs. 0.006 (0.005 – 0.007), IT^PlxnD1^ FLA 0.006 (0.005 – 0.008) vs. 0.006 (0.004 – 0.008), PT^Fezf2^ frontal 0.01 (0.008 – 0.014) vs. 0.008 (0.006 – 0.011), PT^Fezf2^ FLA 0.011 (0.008 – 0.014) vs. 0.007 (0.006 – 0.01)). While no such gross deficits were observed on disrupting IT^PlxnD1^ activity (Fig. 5e,f,g), inhibiting both frontal and frontolateral resulted in a more subtle effect where in mice had difficulty in using fingers to grasp the pellet properly, resulting in decreased agility and spending significantly longer time handling the pellet close to the mouth (Fig. 5e, Suppl. Video 10). Indeed, we found that during IT^PlxnD1^ inhibition the hands were closer to the mouth for a significantly longer time than control trials (Fig 5h hand near mouth duration (sec) control vs inhibition median (1^st^ – 3^rd^ quartile), frontal 0.94 (0.69 – 1.28) vs. 1.14 (0.83 – 1.52), FLA 0.88 (0.61 – 1.27) vs. 1.13 (0.67 – 1.69)). The decreased agility was accompanied by a drop in the number of grasps and rigid finger movements, resulting in a decrease in the distance between fingers during food handling (Suppl. Video 10). To quantify this, we measured the distance between each finger across time (Fig, 5i). We then compared the average inter-finger distance and found it significantly reduced during inhibition of IT^PlxnD1^ in frontal and frontolateral nodes compared to control trials indicating an increase in rigidity of finger movements (Fig 5j, inter finger 1 - finger 2 distance (au) control vs. inhibition median (1^st^ – 3^rd^ quartile), frontal 0.08 (0.075 – 0.087) vs. 0.077 (0.071 – 0.083), FLA 0.082 (0.077 – 0.088) vs. 0.079 (0.073 – 0.084)). These results provide causal evidence that IT^PlxnD1^ and PT^Fezf2^ neurons within the same cortical regions differentially contribute to controlling distinct motor actions of feeding. While PT^Fezf2^ is associated with controlling major oral and forelimb movements including lick and hand lift, IT^PlxnD1^ is likely involved in finer scale coordination such as finger movements during food handling.

**Figure 5.**
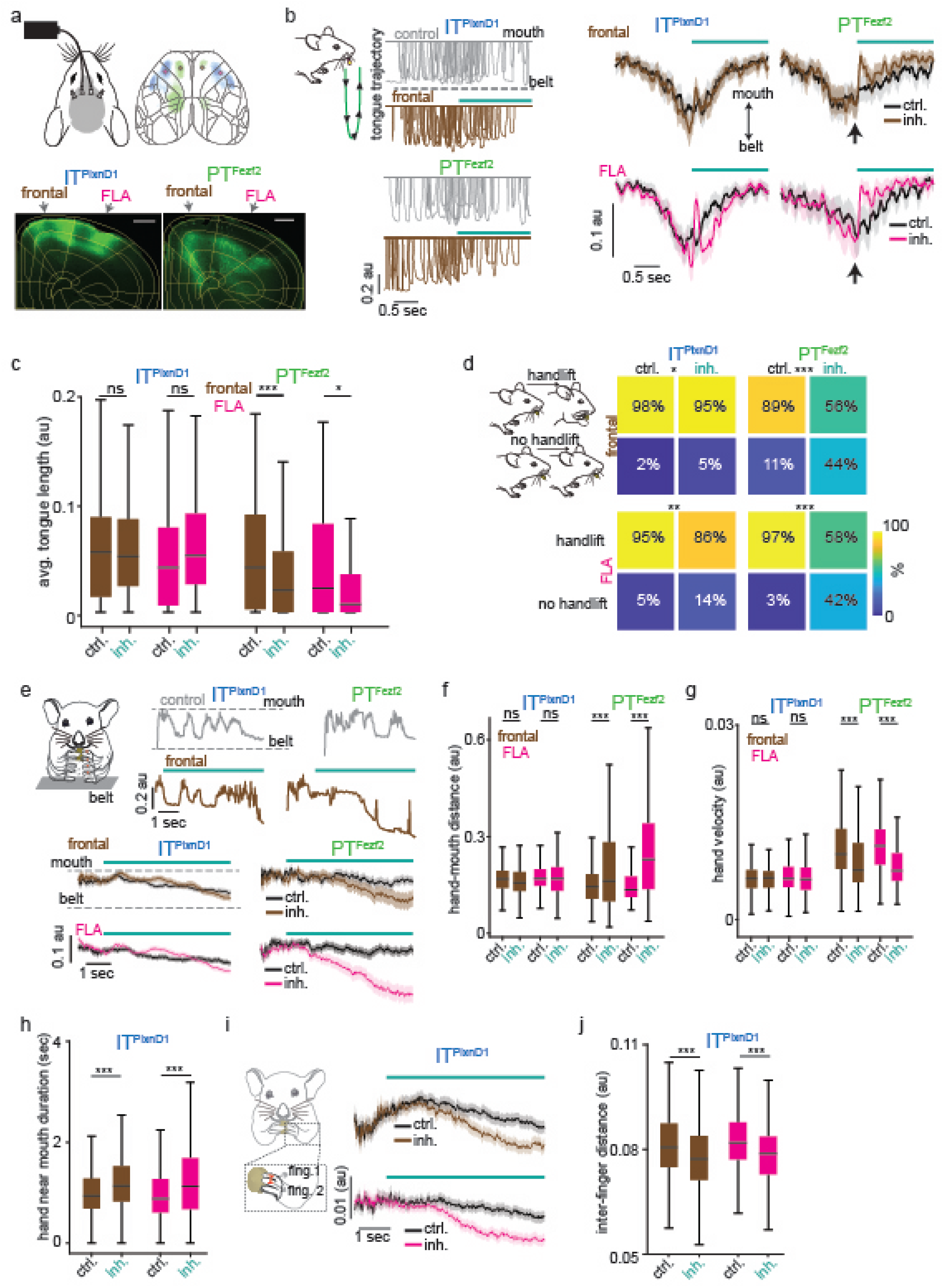
Inhibiting IT^PlxnD1^ and PT^Fezf2^ disrupts distinct components of feeding behavior. a. Top: Schematic of optogenetic inhibition setup (left) with inhibition locations displayed over the Allen dorsal cortex map (right). Brown: Frontal nodes; Magenta: Frontolateral Anterior (FLA) nodes. Bottom: Example coronal sections from IT^PlxnD1^ (left) and PT^Fezf2^ (right) left cortex showing expression of *Gt*ACR1 overlaid by Allen reference map with location of frontal and FLA nodes. White scale bar = 500 μm. b. Left: 10 example tongue trajectories centered to bilateral inhibition of IT^PlxnD1^ (top) and PT^Fezf2^ (bottom) frontal (brown) nodes. Colored trajectories indicate inhibition trials and grey trajectories indicate control trials preceding inhibition trials. Note that the trajectories proceed from the mouth (top) to the pellet (bottom). Green schematic illustrates the progression of a single lick trajectory. Right: Mean tongue trajectories centered to bilateral inhibition of IT^PlxnD1^ frontal (top left), IT^PlxnD1^ FLA (magenta, bottom left), PT^Fezf2^ frontal (top right) and PT^Fezf2^ FLA (bottom right) nodes. Colored trajectories indicate inhibition trials and black trajectories indicate control trials. An upward change in mean value indicates a decrease in tongue trajectory. IT^PlxnD1^ frontal: 173 trials from 4 mice, IT^PlxnD1^ FLA: 165 trials from 4 mice, PT^Fezf2^ frontal: 140 trials from 3 mice, PT^Fezf2^ FLA: 98 trials from 3 mice. c. Distribution of mean tongue length for 1.5 seconds during inhibition and control trials for IT^PlxnD1^ (blue) and PT^Fezf2^ (green) following bilateral inhibition of frontal (brown) and FLA (magenta) nodes (one-way ANOVA (*F_7,1144_=10.02, p=3e-12)* with Tukey-Kramer post hoc multiple comparison test IT^PlxnD1^ frontal control vs. inhibition (*p=0.99),* IT^PlxnD1^ FLA control vs. inhibition (*p=0.29),* PT^Fezf2^ frontal control vs. inhibition (*p=0.002),* PT^Fezf2^ FLA control vs. inhibition (*p=0.02)*). d. Probability of hand lift events during bilateral inhibition of IT^PlxnD1^ frontal (top left), IT^PlxnD1^ FLA (bottom left), PT^Fezf2^ frontal (top right), PT^Fezf2^ FLA (bottom right) nodes (chi-Squared test, IT^PlxnD1^ frontal (*χ=2.89, p=0.05),* IT^PlxnD1^ FLA (*χ=10.75, p=0.0.001),* PT^Fezf2^ frontal (*χ^2^=22.32, p=2.3e-6),* PT^Fezf2^ FLA (*χ^2^=17.05, p=3.6e-5)).* IT^PlxnD1^ frontal: 201 trials from 4 mice, IT^PlxnD1^ FLA: 200 trials from 4 mice, PT^Fezf2^ frontal 82 trials from 3 mice, PT^Fezf2^ FLA 38 trials from 3 mice. e. Top two rows: one example single hand trajectory centered to bilateral inhibition of IT^PlxnD1^ (left) and PT^Fezf2^ (right) frontal (brown) nodes. Colored trajectories indicate inhibition trials and grey trajectories indicate control trials preceding inhibition trials. Note that the trajectories proceed from the belt (bottom) to the mouth (top). Bottom two rows: Mean single hand trajectories centered to bilateral inhibition of IT^PlxnD1^ frontal (top left), IT^PlxnD1^ FLA (bottom left), PT^Fezf2^ frontal (top right) and PT^Fezf2^ FLA (bottom right) nodes. Colored trajectories indicate inhibition trials and black trajectories indicate control trials. A downward change in mean value indicates a deficiency in hand trajectory. IT^PlxnD1^ frontal: 173 trials from 4 mice, IT^PlxnD1^ FLA: 165 trials from 4 mice, PT^Fezf2^ frontal: 140 trials from 3 mice, PT^Fezf2^ FLA: 98 trials from 3 mice. f. Distribution of mean normalized hand to mouth distance for 5 seconds during inhibition and control trials for IT^PlxnD1^ (blue) and PT^Fezf2^ (green) following bilateral inhibition of frontal (brown) and FLA (magenta) nodes (one-way ANOVA (*F_7,2370_=42.48, p=9e-57)* with Tukey-Kramer post hoc multiple comparison test IT^PlxnD1^ frontal control vs. inhibition (*p=0.96),* IT^PlxnD1^ FLA control vs. inhibition (*p=0.99),* PT^Fezf2^ frontal control vs. inhibition (*p=8.4e-8),* PT^Fezf2^ FLA control vs. inhibition (*p=5.9e-8)*). IT^PlxnD1^ frontal: 353 trials from 4 mice, IT^PlxnD1^ FLA: 455 trials from 4 mice, PT^Fezf2^ frontal: 167 trials from 3 mice, PT^Fezf2^ FLA: 202 trials from 3 mice. g. Distribution of mean absolute hand velocity for 5 seconds during inhibition and control trials for IT^PlxnD1^ (blue) and PT^Fezf2^ (green) following bilateral inhibition of frontal (brown) and FLA (magenta) nodes (one-way ANOVA (*F_7,2370_=75.42, p=6e-99)* with Tukey-Kramer post hoc multiple comparison test IT^PlxnD1^ frontal control vs. inhibition (*p=1),* IT^PlxnD1^ FLA control vs. inhibition (*p=0.76),* PT^Fezf2^ frontal control vs. inhibition (*p=0.0001),* PT^Fezf2^ FLA control vs. inhibition (*p=6.1e-8)*). IT^PlxnD1^ frontal: 353 trials from 4 mice, IT^PlxnD1^ FLA: 455 trials from 4 mice, PT^Fezf2^ frontal: 167 trials from 3 mice, PT^Fezf2^ FLA: 202 trials from 3 mice. h. Distribution of mean hand near mouth duration for 5 seconds during inhibition and control trials for IT^PlxnD1^ following bilateral inhibition of frontal (brown) and FLA (magenta) nodes (one-way ANOVA (*F_3,1628_=19.9, p=1.14e-12)* with Tukey-Kramer post hoc multiple comparison test frontal control vs. inhibition (*p=2.7e-5),* FLA control vs. inhibition (*p=8.4e-9).* IT^PlxnD1^ frontal Inh: 353 trials and control: 363 trials from 4 mice, IT^PlxnD1^ FLA Inh: 455 trials and control: 461 trials from 4 mice. i. Mean inter finger 1 - finger 2 distance centered to bilateral inhibition of IT^PlxnD1^ frontal (brown, top) and FLA (magenta, bottom) nodes. Colored trajectories indicate inhibition trials and black trajectories indicate control trials. A downward change in mean value indicates a decrease in inter finger 1 - 2 distance and decrease in finger mobility. Red line over finger 1 and 2 illustrates the variable measured. IT^PlxnD1^ frontal Inh: 353 trials and control: 363 tirals from 4 mice, IT^PlxnD1^ FLA Inh: 455 trials and control: 461 trials from 4 mice. j. Distribution of mean inter finger 1 - 2 distance for 5 seconds during inhibition and control trials for IT^PlxnD1^ following bilateral inhibition of frontal (brown) and FLA (magenta) nodes (one-way ANOVA (*F3,1624=22.85, p=1.72e-14)* with Tukey-Kramer post hoc multiple comparison test frontal control vs. inhibition (*p=2.4e-5),* FLA control vs. inhibition (*p=4e-9).* IT^PlxnD1^ frontal Inh: 353 trials and control: 363 trials from 4 mice, IT^PlxnD1^ FLA Inh: 455 trials and control: 461 trials from 4 mice. *p<0.05, **p<0.005, ***p<0.0005.

### Distinct projection patterns of IT^PlxnD1^ frontolateral and PT^Fezf2^ parietofrontal subnetworks

To explore the anatomical basis of IT^PlxnD1^ and PT^Fezf2^ subnetworks revealed by wide-field calcium imaging, we examined their projection patterns by anterograde tracing using recombinase-dependent AAV in driver mouse lines. Using serial two photon tomography (STP) across the whole mouse brain ^48^, we extracted the brain wide axonal projections and registered them to the Allen mouse Common Coordinate Framework (CCFv3, Methods, ^49,50^). Using 3D masks generated from the CCF-registered map for each brain, we quantified and projected axonal traces within specific regions across multiple planes. With an isocortex mask, we extracted axonal traces specifically within the neocortex and projected signals to the dorsal cortical surface (Fig. 6a).

**Figure 6.**
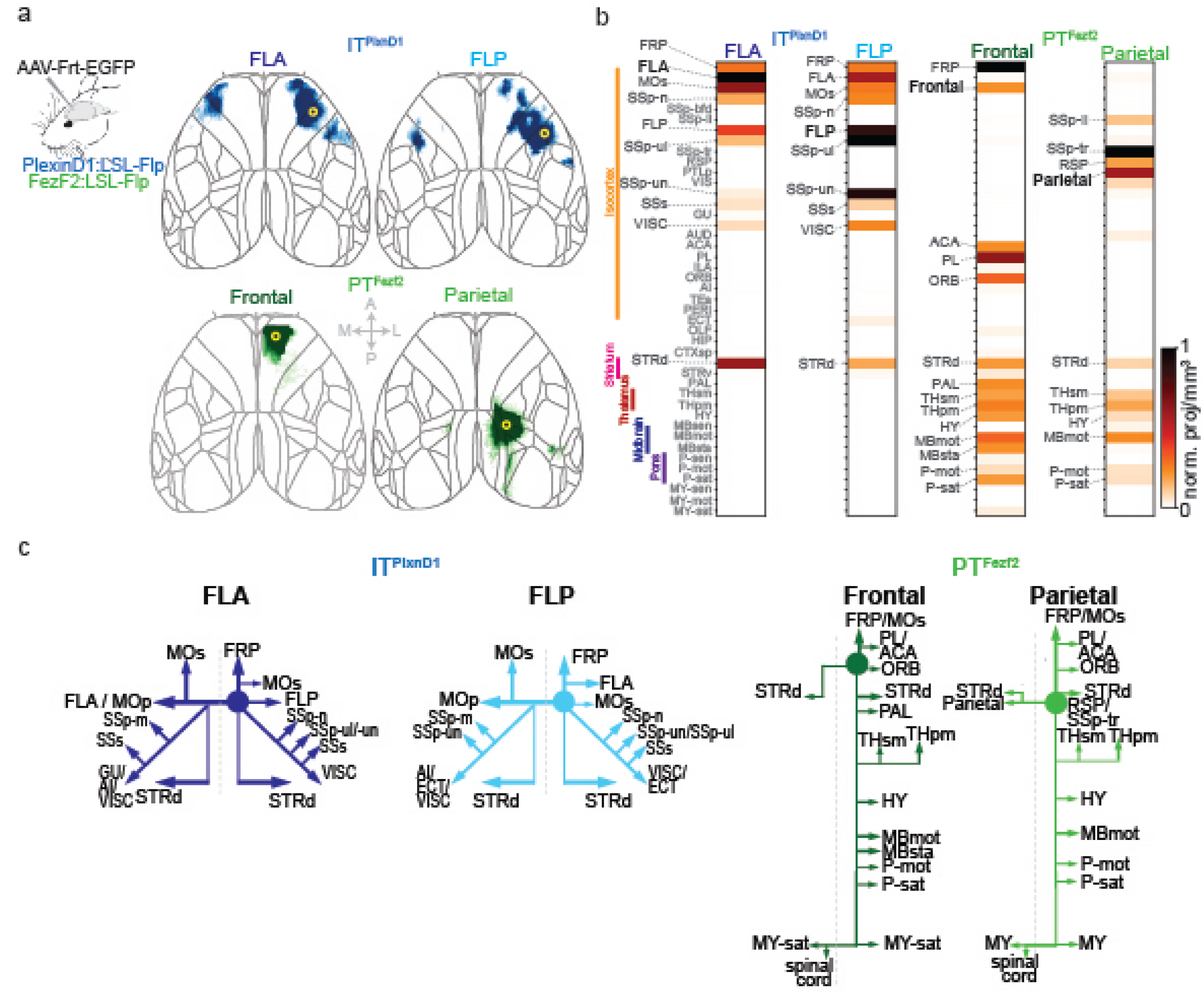
Brain wide projections of IT^PlxnD1^ and PT^Fezf2^ from frontolateral and parietofrontal networks. a. Anterograde projections of IT^PlxnD1^ from FLA and FLP and PT^Fezf2^ from frontal and parietal nodes within isocortex projected to the dorsal cortical surface from an example mouse. b. Brain-wide volume and peak normalized projection intensity maps of IT^PlxnD1^ from FLA and FLP and PT^Fezf2^ from frontal and parietal nodes from an example mouse. Black font indicates injection site; larger gray font indicates regions with significant projections; smaller gray font indicates regions analyzed. c. Schematic of the projection of IT^PlxnD1^ from FLA and FLP and PT^Fezf2^ from frontal and parietal nodes. Circle indicates the site of injection.

As expected, PT^Fezf2^ in parietal and frontal regions show very little intracortical projections (Fig 6a, Extended data 10a, Suppl. Videos 13,14), thus we focused on characterizing their subcortical targets (Fig. 6b, Extended data 10b-e, Suppl. Videos 12,13). PT^Fezf2^ in frontal node predominantly project to dorsal striatum, pallidum (PAL), sensorimotor and polymodal thalamus (THsm, THpm), hypothalamus (HY), motor and behavior state related midbrain regions (MBmot, MBsat), and motor and behavior state related Pons within the hindbrain (P-mot, P-sat, Fig 6b, Extended data 10b-e). PT^Fezf2^ in parietal node projected to a similar set of subcortical regions as those of frontal node, but often at topographically different locations within each target region (Fig. 6b, Extended data 10b-e). To analyze the projection patterns, we projected axonal traces within the 3D masks for each region across its coronal and sagittal plane. For example, PT^Fezf2^ in parietal and frontal nodes both projected to the medial regions of caudate putamen (CP), with frontal node to ventromedial region and parietal node to dorsomedial regions (Extended data 9b); they did not project to ventral striatum. Within the thalamus, frontal PT^Fezf2^ project predominantly to ventromedial regions both in primary and association thalamus while parietal PT^Fezf2^ preferentially targeted dorsolateral regions in both subregions (Extended data 10c). Within the midbrain, we mainly examined projections across collicular regions and found that frontal and parietal PT^Fezf2^ specifically targeted the motor superior colliculus (SCm) with no projections to sensory superior colliculus or inferior colliculus. Within SCm, frontal PT^Fezf2^ preferentially targeted ventrolateral regions while parietal PT^Fezf2^ projected to dorsomedial areas (Extended data 10d). Together, the large set of PT^Fezf2^ subcortical targets may mediate the intention, preparation, and coordinate the execution of tongue and forelimb movements during pellet retrieval and handling. In particular, the thalamic targets of parietofrontal nodes might project back to corresponding cortical regions and support PT^Fezf2^-mediated cortico-thalamic-cortical pathways, including parietal-frontal communications^40^.

In contrast to PT^Fezf2^, IT^PlxnD1^ formed extensive projections within cerebral cortex and striatum (Fig. 6b, Extended data 10b, Suppl. Videos 11,12). Within the dorsal cortex, IT^PlxnD1^ in FLA projected strongly to FLP and to contralateral FLA, and IT^PlxnD1^ in FLP projected strongly to FLA and to contralateral FLP. Therefore, IT^PlxnD1^ mediate reciprocal connections between ipsilateral FLA-FLP and between bilateral homotypic FLA and FLP (Fig. 6a). In addition, IT^PlxnD1^ from FLA predominantly project to FLP (MOp), lateral secondary motor cortex (MOs), forelimb and nose primary sensory cortex (SSp-ul, SSp-n), secondary sensory cortex (SSs) and visceral areas (VISC). Interestingly, IT^PlxnD1^ in FLP also projected to other similar regions targeted by FLA (Fig. 6b). Beyond cortex, IT^PlxnD1^ in FLA and FLP projected strongly to the ventrolateral and mediolateral division of the striatum, respectively (STRd, Fig. 6b, Extended data 10b). These reciprocal connections between FLA and FLP and their projections to other cortical and striatal targets likely contribute to the concerted activation of bilateral FLA-FLP subnetwork during pellet eating bouts involving coordinated mouth-hand sensorimotor actions. Therefore, as driver lines allow integrated physiological and anatomical analysis of the same PN types, our results begin to uncover the anatomical and connectional basis of functional IT^PlxnD1^ and PT^Fezf2^ subnetworks.

### IT^PlxnD1^ and PT^Fezf2^ show distinct spatiotemporal dynamics and spectral properties under ketamine anesthesia

Given the distinct spatiotemporal activation patterns of IT^PlxnD1^ and PT^Fezf2^ in wakeful resting states and sensorimotor behaviors, we further explored whether they differ in network dynamics in a dissociation-like brain state under ketamine/xylazine anesthesia ^37,51^. We first sampled the temporal activity patterns across 6 different regions of dorsal cortex and found that both cell types showed robust oscillatory activity, with IT^PlxnD1^ displaying faster dynamics than PT^Fezf2^ (Extended data 11a, Suppl. Videos 15,16). Quantification of the spectrograms and relative power spectral densities across cortical regions showed that IT^PlxnD1^ exhibited oscillations at approximately 1-1.4 Hz while PT^Fezf2^ fluctuated at 0.6-0.9 Hz (Extended data 11b-d, 12a; IT^PlxnD1^ vs. PT^Fezf2^ 0.6-0.9 Hz. relative power median (1^st^ – 3^rd^ quartile), MOp 0.0035 (0.0025 – 0.0038) vs. 0.003 (0.002 – 0.0042), RSp 0.0022 (0.0016 – 0.0032) vs. 0.0042 (0.0015 – 0.0063), IT^PlxnD1^ vs. PT^Fezf2^ 1-1.4 Hz. relative power median (1^st^ – 3^rd^ quartile), MOp 0.0021 (0.0017 – 0.0029) vs. 0.0009 (0.0006 – 0.0012), RSp 0.0041 (0.0031 – 0.0051) vs. 0.0028 (0.0018 vs. 0.0034)).

To visualize a map of cortical regions strongly oscillating at these frequencies, we computed a map of averaged relative power between 0.6-0.9 and 1-1.4 Hz for each pixel (Extended data 11e, Methods). We then compared the average power distribution at each pixel to identify regions that were significantly different between the two populations (Extended data 12b). While IT^PlxnD1^ was strongly active within the frontolateral at both frequency bands, PT^Fezf2^ was predominantly active in the retrosplenial regions at 0.6 −0.9 Hz (Extended data 11e,12b). To determine distinct pattern of power distribution across mice and sessions, we performed PCA on these spatial maps and projected them onto the top two dimension for each frequency band (Extended data 12c). IT^PlxnD1^ and PT^Fezf2^ both clustered independently with further segregation between IT^PlxnD1^ 0.6-0.9 Hz and 1-1.4 Hz frequency bands (Extended data 12c). Furthermore, these two PN types manifested distinct spatial dynamics across the dorsal cortex, which was visualized by generating a space-time plot displaying activity from pixels obtained across a slice of the cortical surface (Methods, Extended data 11f). While IT^PlxnD1^ displayed complex spatiotemporal patterns with activities spreading, for example, from central regions to anterolateral and posteromedial or from posteromedial to anterolateral and back, PT^Fezf2^ activities predominantly originated from retrosplenial regions and spread to anterolateral regions (Extended data 11f, Suppl. Videos 15,16). To identify the most dominant spatial activation pattern, we performed seqNMF ^52,53^ on the activity data and found that the top dimension accounted for more than 80 % of the variance in IT^PlxnD1^ and over 90% in PT^Fezf2^. Furthermore, we found significant differences in the spatial propagation of activities between the two cell types: whereas IT^PlxnD1^ activities spread multidirections across most of the dorsal cortex, PT^Fezf2^ activities mainly propagated from retrosplenial toward the frontolateral regions (Extended data 11g, 12d). These results show that even under unconstrained brain state such as during anesthesia, IT^PlxnD1^ and PT^Fezf2^ subnetworks operate with distinct spatiotemporal dynamics and spectral properties, likely reflecting their differences in biophysical, physiological ^54^, and connectional properties (e.g. Fig 6).

To examine whether the distinct activity networks observed between IT^PlxnD1^ and PT^Fezf2^ across brain states can simply be explained by their differential cellular density distribution, we obtained GCaMP6f expression for each population using serial two-photon tomography and extracted its mean distribution across the dorsal cortex after registering it to the Allen ccf v3 cortical map (Extended data 13a). We then measured Pearson’s correlation between GCaMP distribution and the activity maps obtained across spontaneous, feeding behavior and ketamine brain states (Extended data 13b). For both populations, we found that the correlations between activity map and GCaMP distribution vary significantly across brain states with values ranging from negative correlations to being close to zero, indicating that cell density distribution patterns alone cannot explain the distinct network dynamics observed between the two cell types (Extended data 13c).

## Discussion

The quest for the cellular basis of cerebral cortex architecture has been fueled by successive waves of technical advances over the past century. Whereas early cytoarchitectonic analyses of cell distribution patterns identified numerous cortical areas ^55^ and characteristic laminar organization across different areas ^56,57^, single cell physiological recording was key to reveal the vertical groupings of neuronal receptive field properties, though initially in anesthetized state ^3,4^. Since its formulation, columnar configuration as the basic units of cortical organization has been a foundational concept that guided decades of research to unravel its cellular and circuitry underpinnings ^5^; yet to date the anatomic basis as well as functional significance of “cortical columns” remain elusive and contentious ^58–60^. Multicellular recordings and computational simulation of neural connectivity led to the hypothesis of a “canonical circuit” template, which may perform similar operations common across cortical areas ^6–8,61^; but the cellular basis of a “canonical circuit” and its relationship to global cortical networks remain unsolved and largely unexplored. An enduring challenge for understanding cortical architecture is its neuronal diversity and wiring complexity across levels and scales ^10^. Meeting this challenge requires methods to monitor and interpret neural activity patterns across multiple layers and large swaths of cortical territory yet with cell type resolution in real time and in behaving animals. Widefield calcium imaging in rodent cortex provide an opportunity to bridge cellular and cortex-wide measurement of neural activity ^22^. Numerous studied have used widefield imaging to investigate cortex wide network dynamics across brain states and behaviors ^26–33^, but most if not all previous studies examined mixed population (e.g. those defined by Thy1 and Rbp4 transgenic lines) comprising different projection classes ^35–38,62^ and thus have yet to compare distinct PN projection types and link anatomical connectivity with functional dynamics. We have recently established genetic tools for dissecting the hierarchical organization PN subpopulations that enable accessing biologically significant PN subpopulations through their inherent developmental, anatomical and physiological properties^39^. Among these, IT and PT represent two major top-level PN classes that mediate intracortical processing and subcortical output channels, respectively, with distinct gene expression ^12,63^, developmental trajectories ^64^, morphological and connectivity features ^41,54^, biophysical properties ^54^, and functional specializations in specific cortical areas^65,66^ and behavior ^67–69^. Several studies have explored the role of IT and PT neurons within individual cortical areas with some suggesting similar tuning properties and others suggesting varying roles depending on behavior and context ^27,65,70–74^, but little is known about their dynamics across global cortical networks. Combining genetic targeting with cortex-wide imaging of their spatiotemporal dynamics, here we demonstrate that IT^PlxnD1^ and PT^Fezf2^ operate in separate and partially parallel subnetworks during a range of brain states and sensorimotor behaviors, and control distinct aspects of feeding movements. These results suggest a revision of the conception of cortical architecture predominantly shaped by the concept of columnar organization ^59^; they indicate that dynamic areal and PN type-specific subnetworks are a key feature of cortical functional architecture that integrates microcircuit components and global brain networks. It is possible that columnar information flow between IT and PT, and thus the functional integration of corresponding subnetworks, might be dynamically gated by inhibitory and modulatory mechanisms according to brain states and behavioral demand.

Modeling and experimental studies have suggested that the source of the signals measured by widefield imaging from cortical surface differs depending on the depth of cell body layer ^75^ and is a weighted average of fluorescence originating across the cortical depth^75,76^. While a large proportion of signal originates from extra somatic layers, especially for deep layer neurons, a significant amount also arises from the cell body layer^75^. Additionally, high correlation between calcium dynamics in cell body and apical dendrites suggest that dendritic signals closely reflect cell body dynamics^42–46^. Furthermore, GCaMP widefield signals are strongly associated with neuronal action potentials both at single cell resolution ^77^ and across cortical depth in a local region ^38,78^. Along with these limitations, it is important to note that GCaMP6f signals have relatively slow temporal dynamics (hundreds of milliseconds); complementary methods with better temporal resolution for spiking activities (e.g. targeted electrophysiological recordings ^79^) are necessary to decipher information flow and neural circuit operation.

Studies in primates and rodents have shown that the posterior parietal cortex (PPC) is a major associational hub that receives inputs from virtually all sensory modalities and frontal motor areas, and supports a variety of functions including sensorimotor transformation, decision making, and movement planning ^80–82^. In particular, PPC subdivisions are strongly connected with frontal secondary motor cortex in a topographically organized manner ^83,84^, and this reciprocally connected network has been implicated in movement intension, planning, and the conversion of sensory information to motor commands ^85^. The cellular basis of parietal-frontal network is not well understood ^81,86^. Here we found that sequential and co-activation of parietal-frontal PT^Fezf2^ neurons are the most prominent and prevalent activity signatures that precede and correlate with tongue, forelimb, and other body part movements. During the hand-assisted pellet feeding task, training mice to eat without hands specifically occluded PT^Fezf2^ parietal activation that normally precedes hand lift. Furthermore, optogenetic inhibition of PT^Fezf2^ within the frontal node disrupts licking and hand lift, while inhibition of the parietal node disrupts hand-to-mouth movement trajectory. Together, these results suggest PT^Fezf2^ as a key component of the parietal-frontal network implicated in sensorimotor transformation and action control. As PT^Fezf2^ neurons do not extend significant intracortical projections, their co-activation in the parietal-frontal network might result from coordinated presynaptic inputs from, for example, a set of IT PNs that communicate between the two areas, or from cortico-thalamic-cortical pathways ^40,87,88^ linking these two areas. Leveraging genetic tools, future PN type-based anterograde, retrograde, and trans-synaptic tracing will reveal the cell type basis of parietal-frontal network. Furthermore, as the topographic connections between parietal and frontal subdivisions appear to correlate with multiple sensory modalities and body axis ^83,84,86^, cellular resolution analysis using two-photon imaging and optogenetically targeted recordings may resolve these topographically organized circuits that mediate different forms of sensorimotor transformation and action control.

While IT^PlxnD1^ neurons show broad and complex activity patterns during several brain states and numerous episodes of sensorimotor behaviors, we discovered a prominent FLP-FLA subnetwork that correlates to a specific aspect of eating involving coordinated mouth and hand movements. Notably, this subnetwork is weakly correlated with pellet retrieval and initial hand lift to mouth, when PT^Fezf2^ in the parietal-frontal subnetwork show strong activation. While FLP mostly comprises primary sensory areas of the orofacial and forelimb regions, FLA comprises frontolateral regions of primary and secondary motor areas. The prominent reciprocal IT^PlxnD1^ projections between these two areas and across bilateral FLP-FLA suggest a compelling anatomical basis underlying the concerted activity dynamics of this functional subnetwork. Furthermore, optogenetic inhibition of IT^PlxnD1^ within frontolateral nodes resulted in finger movement deficits during pellet handling. Together, these results suggest a significant role of the FLP-FLA subnetwork in the sensorimotor coordination of orofacial and forelimb movement during feeding. Systematic cell type-based studies of FLP and FLA may reveal the neural circuit basis underlying oro-manual dexterity for food handling and eating behaviors.

Given the vast diversity of PN types, our focus on IT^PlxnD1^ and PT^Fezf2^ populations in the current study does not achieve a full description of cortical network operations. Indeed, top-level classes further include cortico-thalamic, near-projecting, and layer 6b populations ^12^; and the IT class alone comprises diverse transcriptomic ^12^ and projection ^17^ types that mediate myriad cortical processing streams ^89^. Although IT^PlxnD1^ represents a major subset, other IT subpopulations remain to be recognized and analyzed using similar approaches. It is possible, for example, that another IT type might feature a direct presynaptic connection to PT^Fezf2^ (e.g. ^7,41^) and share a more similar spatiotemporal activity pattern and closer relationship to the PT^Fezf2^ subnetwork. Future studies will leverage emerging genetic tools to more systematically examine additional sets of PN types and subpopulations, thereby achieving an increasingly more comprehensive view of functional cortical networks. The proper granularity of cell type targeting need to depend on the question being addressed and match the granularity of behavior and functional readout. Furthermore, simultaneous analysis of two or more PN types in the same animal will be particularly informative in revealing their functional interactions underlying cortical processing.

## Acknowledgments

We thank Joshua Hatfield for performing STP imaging of PN projections. We thank Bor-Shuen Wang for performing two-photon imaging experiments. We are grateful to Anne Churchland for the numerous discussions, and to Tatiana Engel and Yanliang Shi for discussions on data analysis. We thank Steve Lisberger, Lindsey Glickfeld for comments on the manuscript. This research was supported NIH grant U19MH114823-01 to Z.J.H. Z.J.H is also supported by a NIH Director’s Pioneer Award 1DP1MH129954-01.

## Contributions

Z.J.H. and H.M. conceived the project. H.M. built setups, performed experiments and analyzed data. Z.J.H. supervised the research. X.A. provided advice on the design of feeding behavior and on building feeding behavior setup, and provided mice. X.H.X performed cell type inhibition experiments. P.M. provided advice for analyzing neural data. H.K. performed STP viral injections and surgeries. S.Z. performed in situ imaging experiments. S.M. offered advice on building wide-field imaging setup. Z.J.H. and H.M wrote the manuscript.

## Data Availability

All data will be made available upon request.

## Code Availability

All code will be made available upon request.

## Note

While this manuscript was in preparation, the following preprint was posted in bioRxiv:

Musall, S., Sun, X.R., Mohan, H., An, X., Gluf, S., Drewes, R., Osten, P., Churchland, A.K. (2021) Pyramidal cell types drive functionally distinct cortical activity patterns during decision-making. https://www.biorxiv.org/content/10.1101/2021.09.27.461599v2.abstract

## Methods

### Animals

57 male and female mice were included as part of the study. To express GCaMP6f within specific projection neuron (PN) population, 14 *FezF2-CreER* and 16 *PlexinD1-CreER* knockin mouse lines generated in the lab were crossed with *Ai148* (The Jackson Laboratory, Strain #030328), a GCaMP6f reporter line. 3 *VGAT-ChR2-EYFP* (The Jackson Laboratory, Strain #014548) that express the blue light activated opsin ChR2 in GABAergic interneuron population were used for optogenetic manipulation. 6 *PlexinD1-CreER* and 4 *FezF2-CreER* crossed with a reporter line expressing LSL-Flp were used for viral expression of flp depended anterograde tracing. 4 *PlexinD1-CreER* and 3 *FezF2-CreER* mice were used for cell type specific inhibition experiments. 4 *PlexinD1-CreER* and 3 *FezF2-CreER* mice crossed with *Ai148* were used for two photon imaging experiments. Expression of reporters were controlled via the intraperitoneal injection of tamoxifen (20mg/ml, dissolved in corn oil) between 1 to 2 months postnatal. All mouse colonies at Cold Spring Harbor Laboratory (CSHL) were maintained in accordance with husbandry protocols approved by the IACUC (Institutional Animal Care and Use Committee) and housed by gender in groups of 2 – 4 with access to food and water *ad libitum* and 12 hour light-dark cycle.

### Surgical procedures

For widefield calcium imaging and optogenetic manipulation, adult mice older than 6 weeks were anesthetized by inhalation of isoflurane maintained between 1-2%. Ketoprofen (5 mgkg–1) was administered intraperitonially as analgesia before and after surgery, and lidocaine (2–4 mg kg-1) was applied subcutaneously under the scalp prior to surgery. Mice were mounted on a stereotaxic headframe (Kopf Instruments, 940 series or Leica Biosystems, Angle Two). An incision was made over the scalp to expose the dorsal surface of the skull and the skin pushed aside and fixed in position with tissue adhesive (Vetbond 3M). The surface was cleared using saline and an outer wall was created using dental cement (C&B Metabond, Parkell; Ortho-Jet, Lang Dental) keeping most of the skull exposed. A custom designed circular head plate was implanted using the dental cement to hold it in place. After cleaning the exposed skull thoroughly, a layer of cyanoacrylate (Zap-A-Gap CA+, Pacer Technology) was applied to clear the bone and provide a smooth surface to image calcium activity or for optogenetic stimulation ^24^. For viral injections, we followed the same anesthesia procedure. Under anesthesia, an incision was made over the scalp, a small burr hole drilled in the skull and brain surface was exposed. A pulled glass pipette tip of 20–30 μm containing the viral suspension was lowered into the brain; a 300–400 nl volume was delivered at a rate of 30 nl min^-1^ using a Picospritzer (General Valve Corp); the pipette remained in place for 10 min preventing backflow, prior to retraction, after which the incision was closed with 5/0 nylon suture thread (Ethilon Nylon Suture, Ethicon) or Tissueglue (3M Vetbond), and mice were kept warm on a heating pad until complete recovery ^39^. For cell type specific optogenetic manipulations, we first drilled through the skull using a 0.5 mm bur bilaterally over the frontal and frontolateral anterior areas in each mouse followed by viral injection (*Gt*ACR1) as described earlier. We then implanted Fiber optic cannulae (outer diameter 1.25 mm ceramic ferrule, 400 μm core, 0.39 NA, R-FOC-L400C-39A, RWD) placing them on surface of the brain without penetrating into tissue and sealed them to the skull using dental cement (Tetric EvoFlow, Ivoclar Vivadent AG) followed by head bar implantation.

### Viruses

For cell type specific anterograde tracing we injected 300-400nl of flp dependent viral tracer (AAV2/8-Cag-fDIO TVAeGFP, UNC Vector Core) in *FezF2-CreER;LSL-Flp* mice at either the frontal node (1.7-1.85mm Anterior, 0.7mm lateral, 1.25 mm ventral) or the parietal node (−1.79 to - 1.91 mm posterior, 1.25 to 1.35 mm lateral, 0.3-0.7 mm ventral) and in *PlexinD1-CreER;LSL-Flp* mice at either FLA (1.7 mm anterior, 2.25 mm lateral, 0.3-0.8 mm ventral) or FLP (0.3mm anterior, 3mm lateral, 0.4-0.8mm dorsal). For cell type specific optogenetic manipulation we injected ~400 nl of cre dependent *Gt*ACR1 (AAVDJ-Cbh-DIO-*Gt*ACR1-eYFP) bilaterally in both frontal and frontolateral nodes in each mouse between 300 – 800 μm deep. Mice were between 7 to 12 weeks during viral injection.

### Whole-brain STP tomography and image analysis

Whole brain STP imaging was performed as described earlier ^48^. Briefly, perfused and post fixed brains of adult mice were embedded in 4% oxidized-agarose in 0.05 M PB, cross-linked in 0.2% sodium borohydrate solution (in 0.05 M sodium borate buffer, pH 9.0–9.5).The entire brain was imaged in coronal sections with a 20× Olympus XLUMPLFLN20XW lens (NA 1.0) on a TissueCyte 1000 (Tissuevision) with a Chameleon Ultrafast-2 Ti:Sapphire laser (Coherent) exciting EGFP at 910 nm. Whole-brain image sets were acquired at 0.875 μm x 0.875 μm sampled for 230-270 z sections with a 50 μm z step size. Images were collected by two PMTs (PMT, Hamamatsu, R3896), for signal and autofluorescent background, using a 560-nm dichroic mirror (Chroma, T560LPXR) and bandpass filters (Semrock FF01-680/SP-25). The image tiles were corrected to remove illumination artifacts along the edges and stitched as a grid sequence. Image processing was completed using ImageJ/FIJI and Adobe/Photoshop software with linear level and nonlinear curve adjustments applied only to entire images.

To extract axonal traces, we first used brainreg ^50 49^ to register whole brain data to 10 μm CCFv3 Allen map ^90^. After registration, whole brain background images were linearly fit to the imaging channel and regressed out. Background corrected signals were z-scored and a threshold ranging 0.5 to 1.5 was manually set for each brain above which signal was binarized. Regions with noise in each frame was manually removed using ImageJ/FIJI. To extract volume and peak normalized projection maps (Fig. 6b), we subdivided the whole brain into 43 major nodes as classified by Allen map tree and grouped all regions below that node as part of the parent node mask. We then extracted total axonal count within each roi mask using the binarized data, scaled by the total volume of each mask and normalized by peak value across all nodes. To obtain spatial distribution of axons, 3D masks from registered Allen map for each node was used to extract signal enclosed within the roi from binarized data. Average of the projections within mask either along the coronal or sagittal dimension was obtained to extract axonal distribution across each plane. Masks were also projected along these dimensions and the edges used to obtain the borders for each node (Extended data 8b-e). All STP imaging analysis was performed on python 3.

### In Situ Hybridization

HCR in situ were performed as described^91^. Probes were ordered from Molecular Instruments. Mouse brain was sliced into 50 μm thick slices after PFA perfusion fixation and sucrose protection. Hybridization chain reaction in situ was performed via free floating method in 24 well plate. First, brain slices were exposed to probe hybridization buffer with HCR Probe Set at 37°C for 24 hours. Brain slices were washed with probe wash buffer, incubated with amplification buffer and amplified at 25°C for 24 hours. On day 3, brain slices were washed, counter stained with DAPI and mounted. PlexinD1 (546 nm), Fezf2 (546 nm) and Satb2 (647 nm) probes were used to examine overlaps between these markers.

### Feeding behavior paradigm

We developed a novel behavior paradigm where in mice use an ethological behavioral sequence to capture, handle and feed on food pellets while being head fixed. 2 seconds after trial start, a food pellet (14, 20 or 45 mg, Dustless Precision pellet, F05684, F0071, F0021, Bio-Serv) is delivered on to a custom designed conveyer belt (B375-150XL, ServoCity) with a small pellet holder (made of Velcro strap) from an automatic food pellet dispenser (80209-45, Lafayette Instruments) after which a servo motor (38335S, ServoCity) maneuvers the conveyer belt to bring pellets close to the head fixed mouse. The mouse uses its tongue to pick food pellet to its mouth; lift its hand from a wooden shaft to hold, handle and bite a part of it; bring hands down clasping the remaining pellet; chew; swallow and bring the clasped pellet back towards mouth to continue feeding. At the end of 15 seconds after trial start, the conveyer belt rotates back to dispose of any pellet that was not picked and then moves back to starting position under the pellet dispenser. Layout of task associated temporal events with probability distribution of lick bout onset, pellet in mouth onset, handlift onset, first handling and chew event times are visualized (Extended data 5a).

We began to train mice at least two days after head bar implantation. All mice were older than 6 weeks when training was initiated. Mice were first food restricted so that they reach about 80 to 85% of their initial body weight with ad libitum access to water. During food restriction, mice are fed food pellets (those used in the task) weighing about 10 to 15% of their body weight everyday. During this period mice were habituated to the setup by head fixing them in increasing durations starting from 10 minutes to 30 minutes over 4 days. After habituation and reaching about 85% of their initial weight, we manually placed food pellets close to its mouth while being head fixed. In the beginning mice pick food with their tongue and try to chew without lifting hands. After a few trials, they begin to lift hands to hold the pellet after picking it with their tongue often within the same session or sometimes after a couple of sessions. Once mice become comfortable in lifting their hands after picking pellet, we begin to deliver the pellet close to mouse automatically by moving the conveyer belt from the pellet dispenser. Most mice learn to perform the task within a week of initiating training. In each session mice consumed about 20 to 80 pellets depending on their body weight and days from onset of training. As days progress, mice tend to eat more pellets. At the end of every session, irrespective of the quantity they consume during the task, we provide food pellets weighing about 10 to 15% of their body weight. We constantly measure the mouse weight and make sure they are always above 85% of their initial weight. As sessions progress, they perform increasing number of trails even if their weight is at 90 % of initial body weight.

### Behavior tracking and classification

Using two high speed cameras (FL3-U3-13S2C-CS, Teledyne FLIR) fitted with varifocal lens (#COT10Z0513CS, B&H), we recorded behavior from both the front and left side of the mouse at 100 frames per second as they performed the task under IR illumination. We used DeepLabCut ^47^ to track a range of task components and body parts from both angles including the pellet, pellet holder, left wrist, lower lip, upperlip, nose, tongue tip, left three fingers and right three fingers (from front view). We developed custom algorithms that use these tracked features to identify and classify different behavior events. To identify onset of lick bouts, we calculated spectrogram of the tongue trajectory and extracted the time varying mean power between 4 to 10 Hz. Using a predefined threshold, we obtained time stamps associated with power rising above threshold and falling below it to obtain the onset and offset times of a lick bout. The onset time of the first lick bout after pellet is delivered to the pellet holder was considered as the lick onset time. The time point when food pellet first crossed the lower lip, after being picked by the tongue, for at least 100 ms was considered as the pellet in mouth onset time. To identify handlift onset, we used left hand trajectory to detect time point when absolute velocity crossed a predefined threshold. To extract episodes of food handling, we built a long short term memory (LSTM) neural network classifier with 100 hidden units using MATLAB neural network tool box. After manually labelling time stamps associated with food handling for a few example trials across mice, we used these binary time stamps as output variable and all z-scored trajectories from front view (including the left three fingers, right three fingers, tongue, pellet, upperlip and lowerlip trajectories) of mice as input variables to train the classifier. We then used this classifier to extract time stamps associated with food handling for the remaining trials. To check for classification accuracy, we manually identified the onset and offset times of food handling episodes in 48 randomly selected behavior videos. We then used the classifier to extract the onset and offset times from these videos and measured the number of events that were correctly classified. About 93% of the handling events were correctly identified. To extract chew episodes, we computed spectrogram of the lower lip trajectory to obtain mean power between 3 to 8 Hz and used 0.05 times the standard deviation of this signal as the threshold above which events were classified as chewing. Only episodes longer than 0.5 seconds were used. We performed similar cross validation as described for the food handling classifier and found that about 83% of the chewing episodes were correctly identified. For optogenetic manipulation experiments, we classified an event as handlift if mice successfully brought either of their hands close to the mouth after picking the pellet. To identify hand position during manipulation, we tracked the location of the first finger (Fig 5e) of one of the hands. Instantaneous hand velocity (speed) was quantified as the absolute value of the first derivative of the hand position with respect to time. Hands were considered close to mouth if the distance between finger and mouth was below a custom defined threshold. Inter finger distance was quantified by extracting the instantaneous Euclidean distance between the position of the 1^st^ and 2^nd^ finger (Fig. 5j). Control trials used for comparison were extracted from a non-inhibition trial just preceding each inhibition trial.

### Wide-field calcium imaging

We used wide field imaging to simultaneously measure GCaMP6f activity across the dorsal cortex. The imaging system used was as described previously ^24^. Briefly, we used a sCMOS (edge 5.5, PCO) camera in combination with a 105 mm focal length top lens (DC-Nikkor, Nikon) and 85 mm (85M-S, Rokinon) bottom lens resulting in a x1.24 magnification and a field of view of about 12.5 x 10.5 mm^2^ with an image resolution of 640 x 540 pixels (about 20 μm per pixel dimension) after 4x spatial binning. Images were acquired at 60 Hz alternating between blue (470 nm collimated LED, M470L3, Thorlabs) and violet (405 nm collimated LED, M405L3, Thorlabs) excitation through the same excitation path with the help of a dichroic mirror (no. 87–063, Edmund optics) between the two paths and a 495 nm long pass dichroic (T495lpxr, Chroma) to project both LEDs onto cortical surface. A 525 nm band-pass filter (no. 86–963, Edmund optics) was placed between the camera and lenses to filter GCaMP emission signal. We used the signal associated with 405 nm excitation to regress out the non-calcium dependent hemodynamic signal and isolate the true calcium dependent signal from the 470 nm excitation as described previously^62^. For each pixel, 405 nm and 470 nm excited signals were first ΔF/F normalized using the median across time series within a trial. Signal associated with 405 nm excitation was filtered with a 330 msec (10 frames) moving average filter and linearly fit to the 470 nm excitation signal to obtain regression coefficients. The raw 405 nm excited signal was scaled using these coefficients and regressed out from the 470 nm excited signal to obtain calcium depended ΔF/F signal. For spontaneous and ketamine anesthetized measurements, since activity was measured for 180 seconds continuously, signal was first detrended by fitting and subtracting a 7^th^ order polynomial to the raw signal associated with 405 nm and 470 nm excitation (Extended data 1c) prior to regressing out non-calcium dependent signal as described before. This resulting imaging rate of 30 frames per second after hemodynamic correction was used for all subsequent analysis of calcium activity. All widefield data were rigidly aligned to the Allen CCFv3 dorsal map using four anatomical landmarks; the left, center and right edges of the anterior ridge between the frontal cortex and the olfactory bulb and the lambda ^24,90^ there by allowing data to be combined across mice and sessions.

### Optogenetic Manipulation

To disrupt cortical activity in VGAT-ChR2 mice during behavior, we built a laser scanning system that can direct laser stimulation unilaterally or bilaterally across the whole dorsal cortex surface. A collimated beam of blue light (470 nm) from a laser (SSL-473-0100-10TN-D-LED, Sanctity Laser) was fed into a 2D galvo system (GVS 002, Thorlabs) that was directed onto cortical surface using custom written software. The system contained an additional path to simultaneously visualize the cortical surface using a camera (BFS-U3-16S2C-CS, TELEDYNE FLIR). Using this system, we directed blue light with a beam diameter of 400 μm (full width half maximum) bilaterally at 30 Hz. Laser power at the stimulation site on the cortical surface was set between 10-15 mW. We bilaterally inhibited cortical areas identified from regions active during the feeding behavior task: FLA (1.6 mm anterior, 2.3 mm lateral), FLP(0.5mm anterior, 3.5 mm lateral), Frontal (2 mm anterior, 1 mm lateral) and Parietal (−1.2 mm posterior, 1.2 mm lateral). Using median onset times associated with lick, pellet in mouth and handlift from previous behavior trials, we turned on the stimulation prior to median lick onset time or during licking prior to median pellet in mouth onset time or after pellet in mouth but prior to median handlift onset time or after hand lift onset time during manipulation, bilaterally inhibiting each of the four ROIs for durations ranging from 5 to 7 seconds. The inhibition was randomly turned on between control trials where we did not provide any laser stimulation. For cell type specific manipulation, we used splitter branching fiber-optic patch cords (400 μm core, SBP(2) 1m FCM-2xZF2.5, doric) attached to the head of a 532 nm laser (GL532T3-100FC, SLOC Lasers). The output fibers were attached bilaterally to either the frontal or frontolateral anterior optic fiber implants during behavior for manipulation of either IT^PlxnD1^ or PT^Fezf2^ neurons. Laser power was set between 5-8 mW. The inhibition protocol was as described earlier.

### Neural and behavior data analysis

All neural and behavior analysis was performed on MATLAB v2018b and Python 3.8/3.9.

#### Wakeful resting state analysis

Mice were first habituated to head fixation in the setup as described earlier. We imaged and extracted calcium dynamics at 30 fps (as described earlier) across the dorsal cortex for 3 minutes from 6 mice for each PN across two sessions on different days. We also simultaneously video recorded behavior at 20 fps. We filtered the calcium activity between 0.01Hz and 5 Hz per pixel to remove the extremely low frequency fluctuations and high frequency noise using 100 order finite impulse response bandpass filter. Power spectral density plots from both PNs during wakeful resting state indicated activity frequency was well below 5 Hz (Extended data 1d). Since most of the calcium activity was well below 5 Hz we also band pass filtered behavior videos between 0.01 and 5 Hz to retain the low frequency components of movements. To obtain time varying behavior variance signal, we computed the differential for each pixel in a video frame across time to obtain motion energy ^92^ and then computed variance for each frame of the motion energy video (Fig. 1c black trace). We resampled these traces to match sampling rate of calcium activity (30 Hz) using resample function in MATLAB. Time points when behavior variance exceeded 0.2 times the standard deviation were classified as active periods and remaining as quiescent. We then split calcium activity videos into active and quiescent episodes aligned with these time stamps and extracted variance per pixel across time for each episode to obtain variance maps. We then peak normalized these maps and averaged them across mice and sessions (Fig. 1d). To compute neural variance explained by spontaneous behavior, we performed singular value decomposition (SVD) of both calcium activity (Cn,t, n: number of pixels, t: number of time points) and motion energy videos (*d*B_b,t_ / *dt*, b: number of pixels, t: number of time points) and extracted the top 200 temporal components (S*V^T^) from each decomposition (Extended data 3b). We then extracted amplitudes of the motion energy temporal components by computing the absolute value of their Hilbert transforms (Extended data 3b). The decomposition of behavior video data using SVD is similar to the method previously described^93^. Here, instead of taking the absolute value of motion energy data prior to applying SVD, we used Hilbert transform to obtain time varying amplitudes on the temporal dimensions obtained after decomposing the motion energy videos (Extended data 3b). This results in obtaining the slow varying dynamics of the motion energy signal that is used to model the relatively low temporal resolution widefield data (Extended data 3b). We then built a liner encoding model using these 200 amplitude signals as independent variables to explain the top 200 temporal components of the calcium activity (Extended data 3b). We performed 5 fold cross validation of the model to obtain the cross validated R^2 92^.

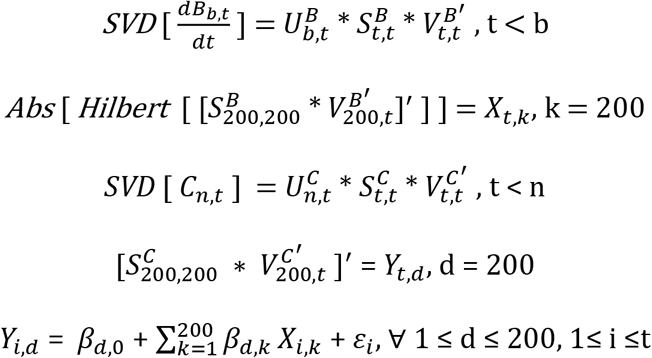

To compute variance explained by each body part, we first defined a window around each body part (Fig. 1f inset) and extracted motion energy within those windows as described earlier (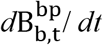, b: number of pixels, t: number of time points, bp: body part). We then extracted amplitudes (absolute of their Hilbert transforms) from each pixel of the motion energy video and averaged them within each window to obtain a time varying amplitude signal associated with movement of each body part (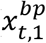, Extended data 3c). We built a linear model using z-scored amplitude signal per body part as independent variables to explain the top 200 SVD temporal components of calcium activity (*Y_t,d_*, t: number of time points, d: number of dimensions). We performed 5 fold cross validation to obtain the cross validated R^2^ associated with each body part. For both linear encoding models we used ridge regression and identified the optimal lambda for each session from a range of values using 5 fold cross validation. To obtain the spatial map of transformed regression weights (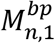, n: number of pixels, bp: body part), we computed the dot product between the model regression weights associated with each body part and the corresponding top 200 SVD spatial components (U) and transformed the resulting vector to a 2D matrix with dimensions matching the calcium activity frame.

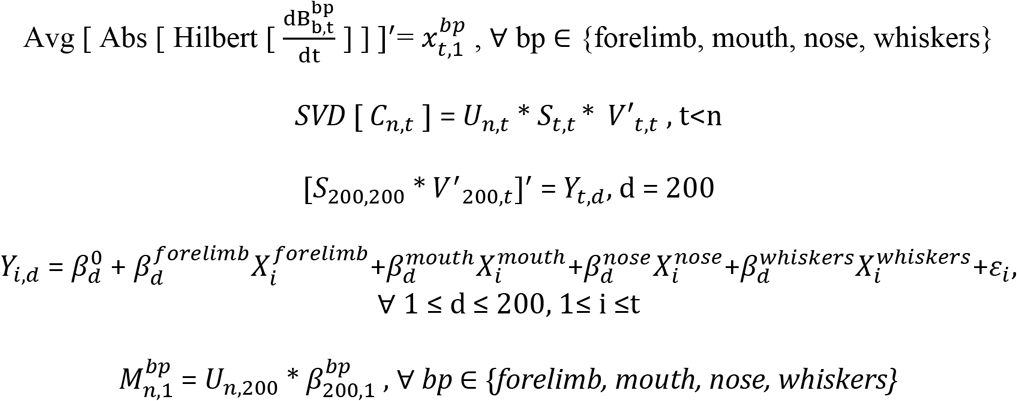

#### Sensory stimulation analysis

Before stimulation, mice were injected with chlorprothixene (1 mg/Kg i.p.) and maintained under light isoflurane anesthesia (0.8-1% with O_2_). We then placed a custom designed cardboard attached to two piezo actuators (BA5010, PiezoDrive) close to left whisker pad between whiskers and just below the upper and lower lip. We also placed an orange LED close to dorsal region of the left eye. We used an Arduino Uno Rev3 (A00006, Arduino) to drive the piezo and LED. A single trial consisted of 3 seconds baseline followed by whisker stimulation at 25 Hz for 1 second, 3 second delay, orofacial stimulation at 25 Hz for 1 second, 3 second delay, blinking visual stimulus at about 16 Hz for 1 second followed by 3 second delay before starting the next trial. We recorded one session per day consisting of 20 trials. To extract temporal traces, we used spatial maps obtained by averaging IT^PlxnD1^ activity per pixel during the 1 second stimulation period in response to each sensory stimulation. We identified centers of peak activity in each map and used a circular window of 560 μm diameter to extract signals within the circular mask and average them per frame.

#### Two photon imaging and analysis

We used a Sutter movable objective microscope to measure single neuron calcium dynamics at 30.9 Hz over the left whisker somatosensory cortex. The location was identified using the peak activity following whisker stimulation from widefield imaging experiments. Each trial consisted of 3 seconds of baseline followed by 1 second whisker stimulation (as described previously) followed by another 3 seconds of post stimulus measurement. For each field of view, we measured responses across 20 trials. We recorded from cell bodies in IT^PlxnD1^ and apical dendrites of PT^Fezf2^ (200 – 500 μm dorsoventral). We did not record from cell bodies in PT^Fezf2^ since they were relatively dim due the depth. Dendritic calcium activity in layer 5B neurons has shown to be strongly correlated to cell body dynamics^42–46^. We used suite2p (https://www.suite2p.org/) to identify neurons and extract calcium dynamics followed by removal of neuropil activity and z-score computation for each neuron. To classify neurons, we used linear modelling to fit the response of each cell to a predictor variable containing ones during whisker stimulation and zeros otherwise. We used statsmodels module in Python to model the fit and obtain regression weights along with the associated statistical significance. Neurons with significantly positive regression weights (p<0.05) were classified as activated while those with significantly negative weights were classified as inhibited neurons. All other neurons were grouped as unclassified.

#### Feeding behavior analysis

To identify sequential activation pattern during feeding behavior, we extracted frames one second before and one second after pellet in mouth onset for all trials across mice and sessions. Since each frame is registered to the Allen CCFv3, we computed mean for each pixel at every sampling point to obtain an average activation map at each time point centered around pellet in mouth.

To identify activation maps associated with specific behavior event, we used a linear modelling approach. We generated binary time stamps associated with each behavior event as independent variables. For a single trial the lick variable consisted of ‘ones’ 0.3 seconds before lick onset to the start of pellet in mouth onset and ‘zeros’ at all other time points. The pellet in mouth variable consisted of ‘ones’ from start of pim onset to hand lift onset and ‘zeros’ everywhere else. The hand lift variable consisted of ‘ones’ from start of hand lift onset to 0.5 seconds after and ‘zeroes’ everywhere else. The handling and chew variables similarly contained ‘ones’ at time stamps classified by the handling and chew classifier previously described and zeros everywhere else. We then concatenated these variables vertically across trials within a session to generate the independent variable matrix (*X_t,5_*, t: number of time points). We then combined calcium activity across trials within a session (*C_n,t_*, n: number of pixels, t: number of time points) and performed SVD to obtain top 200 temporal components (*S*V^T^*) and used a linear model to fit the independent behavior variables to the SVD temporal components obtained from neural activity. We then computed a dot product between the regression weights obtain from the model for each behavior variable and spatial components (U) of the neural activity SVD to transform the regression weights into a spatial map (M) of regions associated with each behavior event.

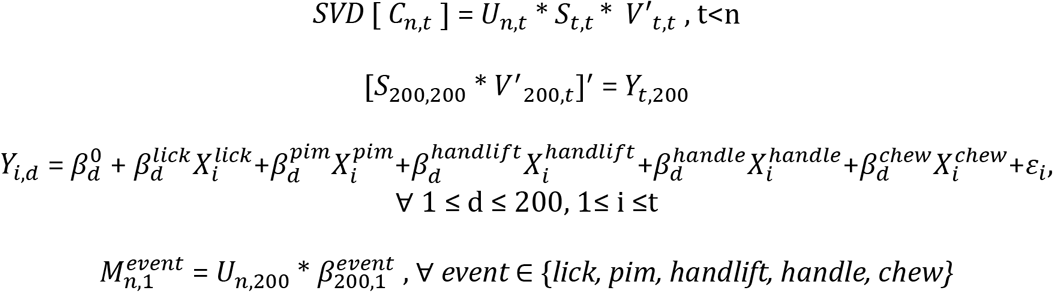

To determine the significance of each independent variable in the model, we first measured the variance explained (R2) from the full model. We then measured the variance explained from a reduced model with time stamps for only one of the variables randomly shuffled and computed the difference in explained variance between the full and reduced model for each variable (delta R2). We computed the distribution of delta R2 for each variable by randomly shuffling time stamps 1000 times per session and measured the difference between full and reduced model and estimated its significance by quantifying the proportion of delta R2 greater than 0 (Extended data 6c). All variables for both PNs significantly contributed to the model with p<0.005 (Extended data 6c).

To identify the center of activation so as to extract temporal traces, we first calculated the average activity per pixel for 1 second before to 2 seconds after pim onset across mice and sessions (Fig. 3a,d). We then applied a mask containing mouth and nose primary sensory dorsal cortex region (as labeled by the Allen CCF V3) over the IT^PlxnD1^ activation map and identified center of peak activation and used it as the center of Frontolateral Posterior node (Fig. 3d, orange). Similarly we used MOs and MOp mask over IT^PlxnD1^ activation map to identify the center of Frontolateral Anterior node (Fig 3d, magenta). We used MOs mask over the PT^Fezf2^ activation map to identify the center of frontal node (Fig 3a, dark brown) and a few cortical regions in the posterior area (RSP_agl_, VIS_am_, VIS_a_, SS_p-tr_, SS_p-ll_, SS_p-ul_, SS_p-un_, ViS_rl_, SS_p-bfd_) to identify the center of parietal node (Fig 3a, light brown). We used a circular window mask of 560 μm diameter around these centers to extract signals within these masks and averaged them per frame to obtain temporal dynamics from each node. The Allen masks were used only to help identify the centers of peak activation and were not used to parcellate the cortex for any analysis.

To identify distinct activation clusters using Linear Discriminant Analysis (LDA), we first combined temporal activity centered to PIM onset from all trials within an ROI from both PNs along the temporal dimension. We then concatenated the PN type class labels associated with each trial and performed LDA on the activity matrix and class labels using the LDA toolbox (LDA: Linear Discriminant Analysis, https://www.mathworks.com/matlabcentral/fileexchange/29673-lda-linear-discriminant-analysis, MATLAB Central File Exchange. Retrieved December 28, 2021). We then projected the temporal activity matrix on the first two dimensions identified by the analysis and colored them based on PN type to visualize clusters.

#### Ketamine anesthetized state analysis

To measure cortical dynamics under anesthesia we injected a mixture of Ketamine (87.5 mgKg^-1^) / Xylazine (12.5 mgKg^-1^) i.p. in each mouse and measure activity for 3 minutes at 30 Hz as described earlier. We used a 560 μm diameter circular mask and extracted average activity per frame within the mask across time to obtain signals from specific regions. We used Chornux ^94^ toolbox to calculate multi-taper spectrograms and relative power spectral densities. To obtain space time plot (Fig. 7f), we extracted pixels across a one dimensional slice of cortical surface and visualized calcium dynamics for each pixel in the slice across time.

To visualize the most dominant spatial activation pattern, we used seqNMF as described previously ^52,53^. Briefly, the calcium activity was first band pass filtered between 0.01 Hz and 5 Hz. Since seqNMF is applied to non-negative matrices, values below zero were set to 0. This 3D matrix is transformed to a 2D matrix of pixels x time and seqNMF with parameters; sequence length: 30 (1 second), number of dimensions: 10 and lambda: 0.01 was used to obtain the most dominant sequence. Across a range of lambda values seqNMF identified just one sequence for both IT^PlxnD1^ and PT^Fezf2^ accounting for greater than 80% of variance. To identify most dominant sequence across mice and sessions, the first 20 seconds of activity from each mouse and session with a PN type was concatenated along the temporal dimension. We then used seqNMF to identify the most dominant sequence for each PN type as described above (Extended data 12d).

---

**Extended Data 1.**
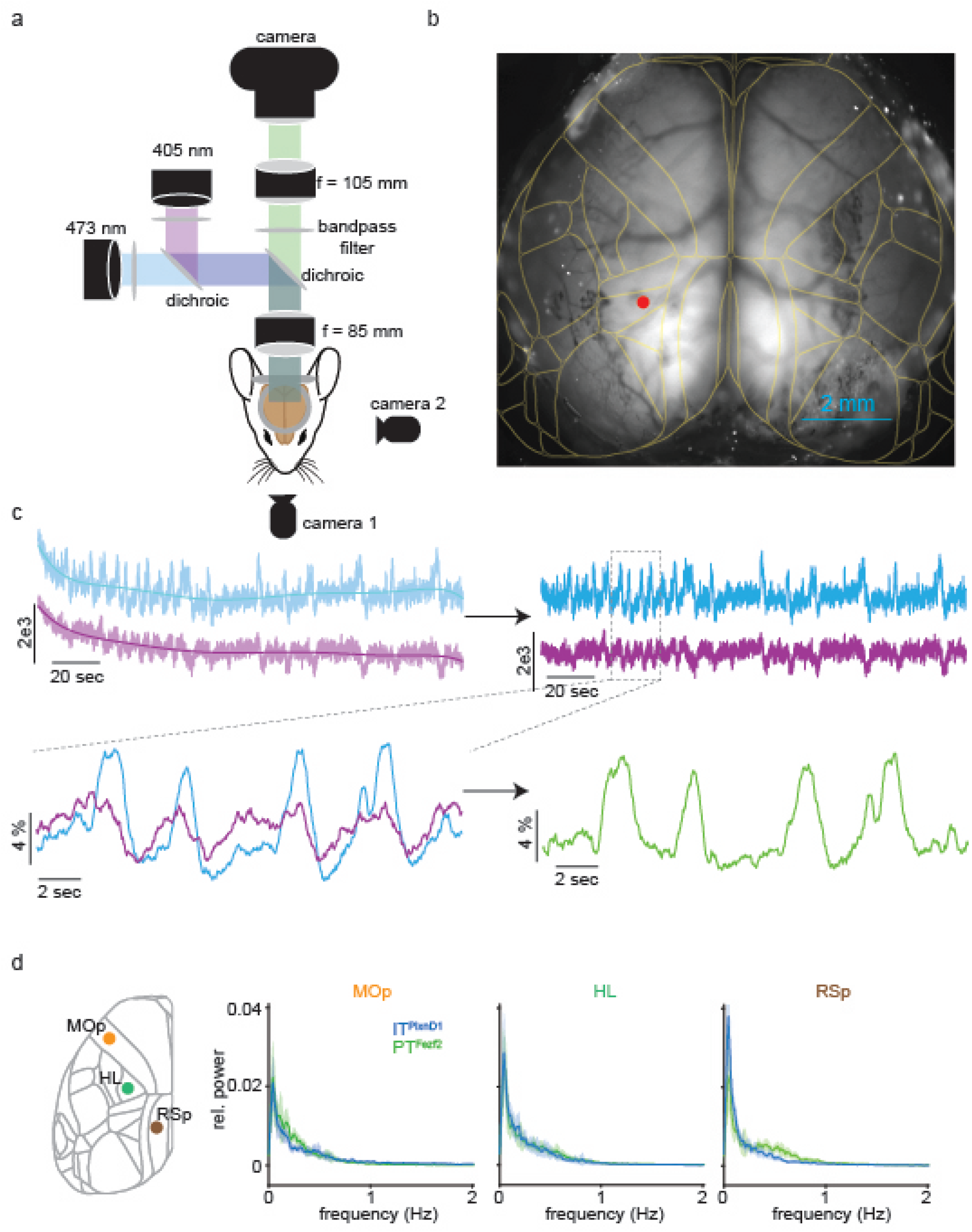
Widefield imaging system layout and signal correction. a. Schematic of the wide-field imaging system. b. Example field of view of dorsal cortex obtained from the widefield imaging system overlaid with registered Allen map in yellow. Red dot indicates the location used to plot example traces in panel c. c. Illustration of the method used to obtain final df/f signal for spontaneous recordings. Blue and violet traces indicate signal obtained during excitation with 473 nm and 405 nm respectively. Top left: raw signal. Top right: signals after detrending. Bottom left: Zoomed in signals after normalizing to baseline (df/f). Bottom right: Final signal after regressing violet from blue trace. d. Mean relative power spectral density of IT^PlxnD1^ (blue) and PT^Fezf2^ (green) activity from MOp, HL sensory and RSp during awake resting state condition (average of 12 sessions across 6 mice, shading around trace ±2 s.e.m).

**Extended Data 2.**
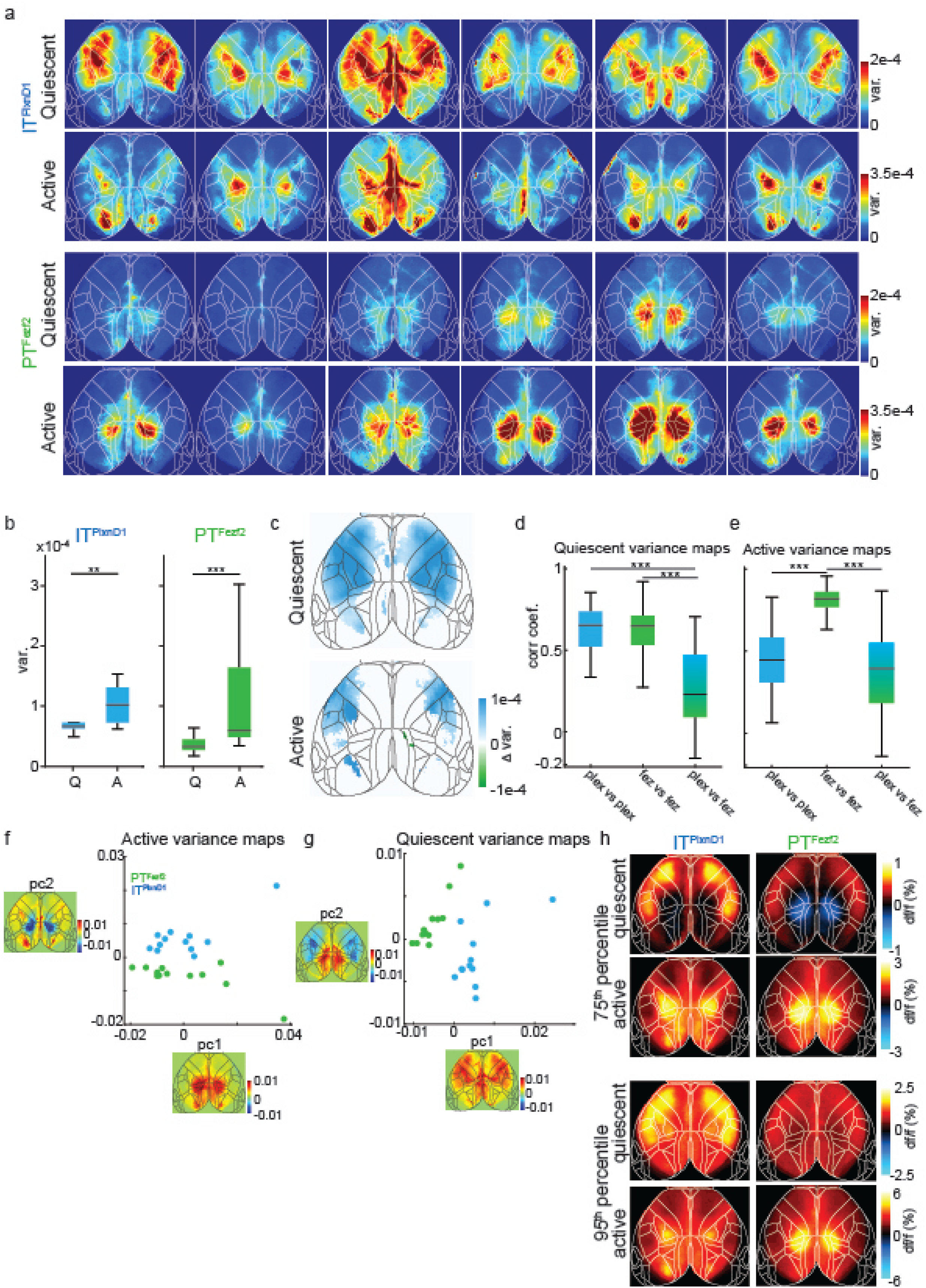
IT^PlxnD1^ and PT^Fezf2^ activation patterns during wakeful resting state across mice. a. Variance maps for each mouse (in columns) during quiescent and active episodes averaged over two sessions. b. Distribution of variance during quiescent (Q) versus active (A) episodes in IT^PlxnD1^ (blue) and PT^Fezf2^ (green) (12 sessions from 6 mice for each cell type), paired two-sided Wilcoxon signed rank test Quiescent vs. Active IT^PlxnD1^ (*U=77, p = 9.8e-4),* PT^Fezf2^ (*U=78, p=4.8e-4)).* c. Difference between IT^PlxnD1^ and PT^Fezf2^ average variance maps for quiescent and active episodes (12 sessions from 6 mice for each cell type). Only significantly different pixels are displayed (*two-sided Wilcoxon rank sum test with p-value adjusted by False Discovery Rate (FDR) = 0.05).* Blue pixels indicate values significantly larger in IT^PlxnD1^ compared to PT^Fezf2^ and vice versa for green pixels. d. Distribution of Pearson’s correlation coefficients between quiescent variance maps within IT^PlxnD1^ (blue), PT^Fezf2^ (green) and between IT^PlxnD1^ & PT^Fezf2^ (blue green) (66 pairs within IT^PlxnD1^ & PT^Fezf2^ and 144 pairs between IT^PlxnD1^ & PT^Fezf2^ in 12 sessions from 6 mice each, Kruskal-Wallis test (*χ^2^(2) = 129.02, p = 9.6e-29)* with Tukey-Kramer (*alpha = 0.05)* post hoc multiple comparison test IT^PlxnD1 vs. PlxnD1^ vs. PT^Fezf2 vs. Fezf2^ (*p=0.86),* IT^PlxnD1 vs. PlxnD1^ vs. IT/PT^PlxnD1 vs. Fezf2^ (*p=5.9e-8)*, PT^Fezf2 vs. Fezf2^ vs. IT/PT^PlxnD1 vs. Fezf2^ (*p=5.9e-8)*). e. Distribution of Pearson’s correlation coefficients between active variance maps within IT^PlxnD1^ (blue), PT^Fezf2^ (green) and between IT^PlxnD1^ and PT^Fezf2^ (blue green) (66 pairs within IT^PlxnD1^ & PT^Fezf2^ and 144 pairs between IT^PlxnD1^ & PT^Fezf2^ in 12 sessions from 6 mice each, Kruskal-Wallis test (*χ^2^(2) = 125.19, p = 6.5e-28)* with Tukey-Kramer (*alpha = 0.05)* post hoc multiple comparison test IT^PlxnD1 vs. PlxnD1^ vs. PT^Fezf2 vs. Fezf2^ (*p=5.9e-8),* IT^PlxnD1 vs. PlxnD1^ vs. IT/PT^PlxnD1 vs. Fezf2^ (*p = 0.32)*, PT^Fezf2 vs. Fezf2^ vs. IT/PT^PlxnD1 vs. Fezf2^ (*p=5.9e-8)*). f. Distribution of IT^PlxnD1^ (blue) and PT^Fezf2^ (green) active variance maps projected to the subspace spanned by the top two principal components. g. Distribution of IT^PlxnD1^ (blue) and PT^Fezf2^ (green) quiescent variance maps projected to the subspace spanned by the top two principal components. h. Average maps of the 75^th^ (top) and 95^th^ (bottom) percentile df/f value during active and quiescent episodes for IT^PlxnD1^ and PT^Fezf2^. *p<0.05, **p<0.005, ***p<0.0005.

**Extended Data 3.**
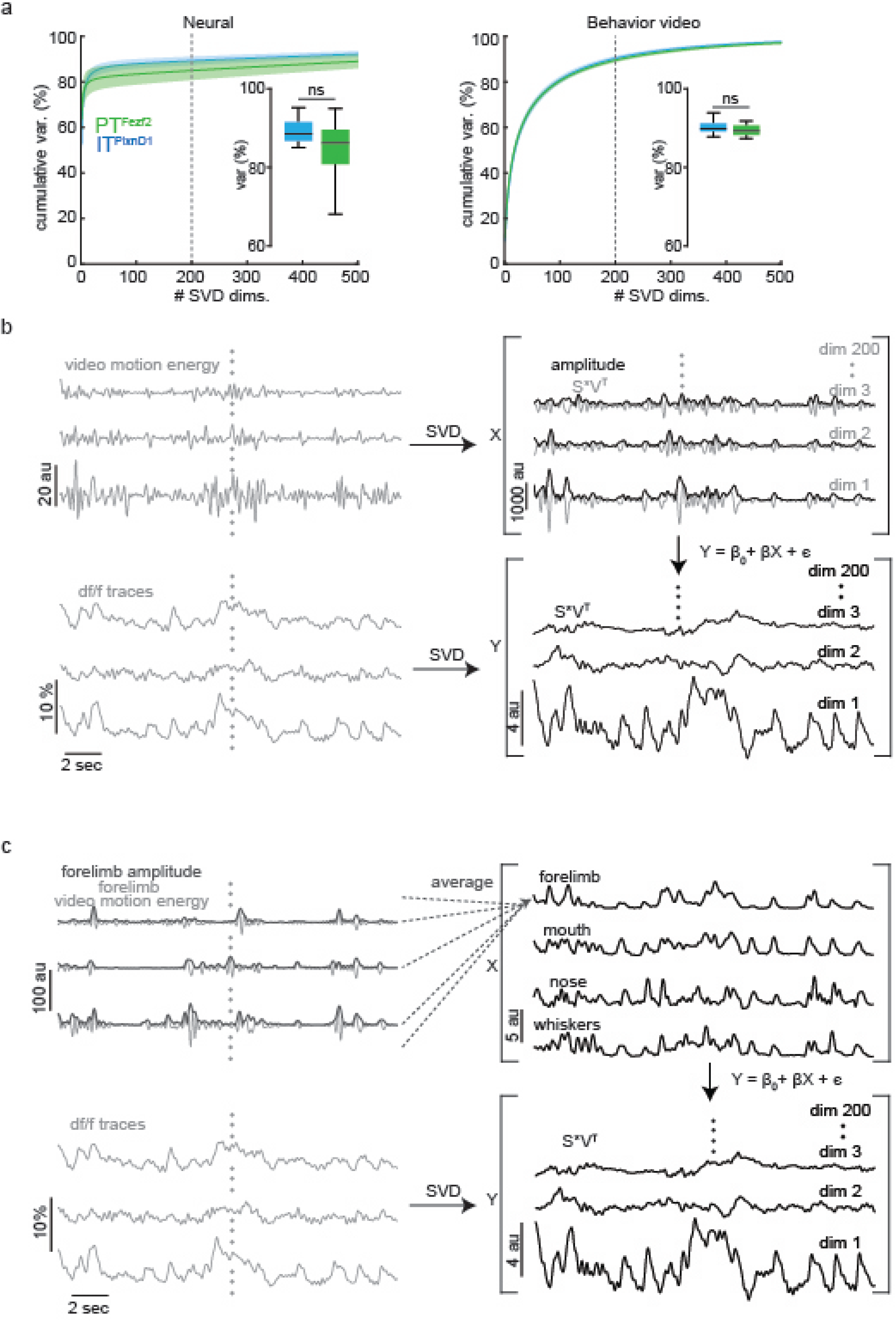
Linear encoding model for measuring neural variance associated with behavior variance. a. Left: Cumulative distribution of variance accounted by the corresponding singular value decomposition (SVD) temporal dimensions of IT^PlxnD1^ (blue) and PT^Fezf2^ (green) calcium activity. Inset: Distribution of IT^PlxnD1^ (blue) and PT^Fezf2^ (green) cumulative variance accounted by the top 200 neural SVD dimensions (two-sided Wilcoxon rank sum test (*U=178, z=1.59, p = 0.11)).* Right: Cumulative distribution of variance accounted by the corresponding SVD temporal dimensions of IT^PlxnD1^ (blue) and PT^Fezf2^ (green) behavior video motion energy (average of 12 sessions across 6 mice for each cell type, shading around trace ±2 s.e.m). Inset: Distribution of IT^PlxnD1^ (blue) and PT^Fezf2^ (green) cumulative variance accounted by the top 200 behavior video SVD dimensions (twosided Wilcoxon rank sum test (*U=168, z=1.01, p = 0.31)).* b. Illustration of the linear encoding model used to fit SVD temporal dimensions of behavior video to neural activity SVD temporal dimensions. Top left: traces indicate motion energy signal of example pixels from behavior video frames. Top right: Top three SVD temporal dimensions along with its amplitudes obtained by decomposing motion energy of behavior videos. Bottom left: calcium traces from example pixels corresponding to the behavior frame. Bottom right: Top three SVD temporal dimensions obtained by decomposing neural activity. c. Illustration of the linear encoding model used to fit behavior variance within specific windows of the behavior video to the top 200 neural activity SVD temporal dimensions. Top left: Traces indicate motion energy signal with its amplitude from example pixels within the forelimb window. Top right: Average of amplitudes from all pixels within the forelimb widow. Bottom left: Calcium traces from exampled pixels corresponding to the behavior frame. Bottom right: Top three SVD temporal dimensions obtained by decomposing neural activity.

**Extended Data 4.**
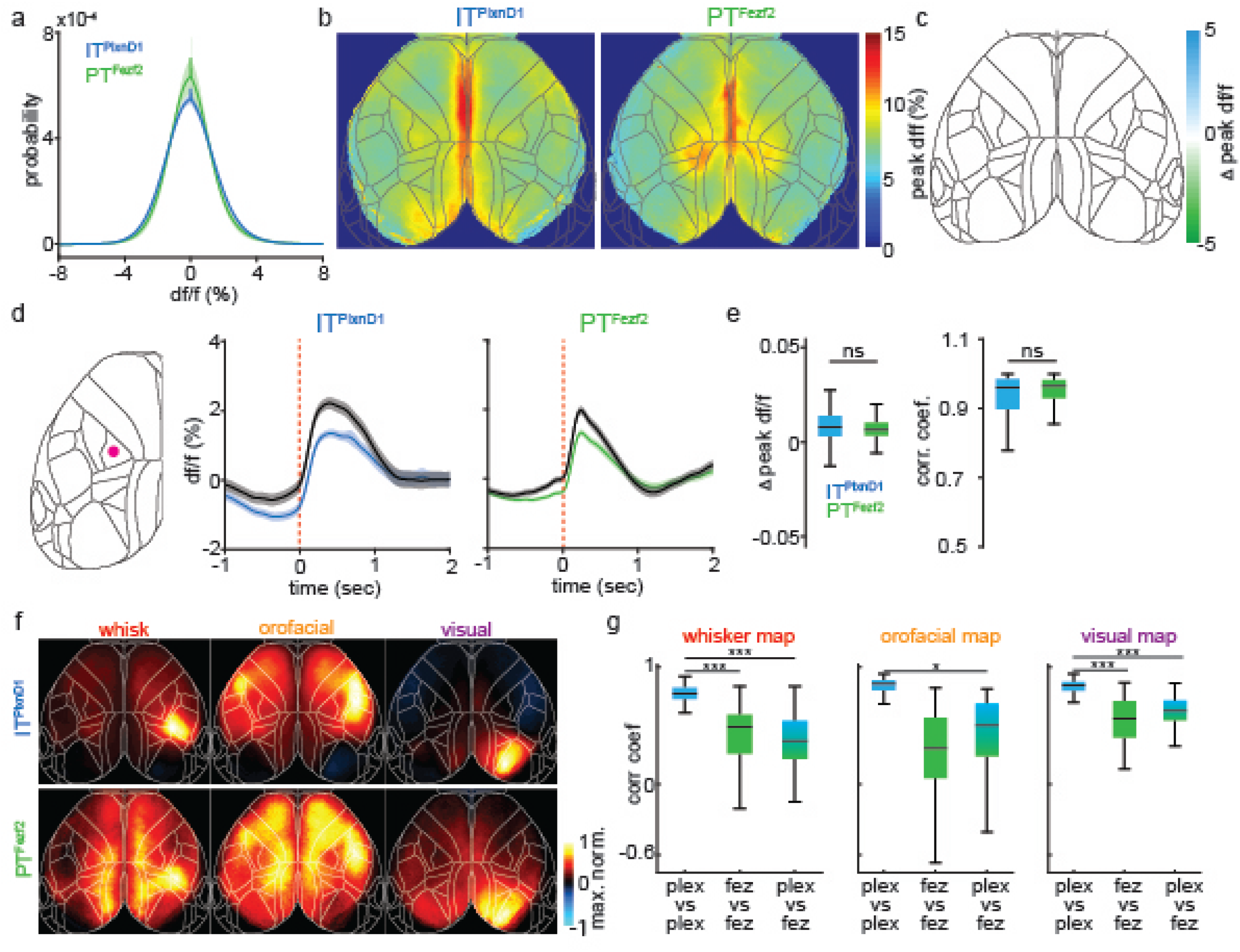
Spontaneous activity comparison and correlation of sensory response in IT^PlxnD1^ and PT^Fezf2^ across mice. a. Probability distribution of df/f values from IT^PlxnD1^ (blue) and PT^Fezf2^ (green) during wakeful resting state (average of 12 sessions from 6 mice each, shading around trace indicate ± 2 s.e. m). b. Mean peak df/f maps of IT^PlxnD1^ (left) and PT^Fezf2^ (right) during spontaneous behavior (average of 12 sessions from 6 mice for each cell type). c. Difference between IT^PlxnD1^ and PT^Fezf2^ mean peak df/f maps (12 sessions from 6 mice for each cell type). Only significantly different pixels are displayed (*two-sided Wilcoxon rank sum test with p-value adjusted by FDR = 0.05).* Note that no pixels are significantly different. d. Mean temporal dynamics of IT^PlxnD1^ (blue) and PT^Fezf2^ (green) activity with (colored) and without (black) hemodynamic correction from hindlimb sensory area during spontaneous behavior. Activity is aligned to the onset of spontaneous movements (average of 367 (IT^PlxnD1^) and 474 (PT^Fezf2^) trials in 12 sessions from 6 mice for each cell type, shaded region indicates ±2 s.e.m). Left image with red dot indicates location used to extract signal. e. Left: Distribution of the difference between hemodynamic corrected and uncorrected peak df/f value between 0 to 1 sec after spontaneous movement onset for IT^PlxnD1^ (blue) and PT^Fezf2^ (green) activity from panel d (367 and 474 trials for IT^PlxnD1^ and PT^Fezf2^, respectively, two sample t-test IT^PlxnD1^ vs. PT^Fezf2^ (*t_839_=1.59, p=0.11)).* Right: Distribution of Pearson’s correlation coefficient between hemodynamic corrected and uncorrected IT^PlxnD1^ (blue) and PT^Fezf2^ (green) activity from panel d (367 and 474 trials for IT^PlxnD1^ and PT^Fezf2^, respectively, two-sided Wilcoxon rank sum test (*U=151312, z=-0.91, p = 0.36)).* f. Mean peak normalized activity maps of IT^PlxnD1^ (top) and PT^Fezf2^ (bottom) in response to corresponding unimodal sensory simulation (average of 12 sessions from 6 mice for each cell type). g. Distribution of Pearson’s correlation coefficients between sensory activation maps within IT^PlxnD1^ (blue), PT^Fezf2^ (green) and between IT^PlxnD1^ and PT^Fezf2^ (blue green) (66 pairs within IT^PlxnD1^ & PT^Fezf2^ and 144 pairs between IT^PlxnD1^ & PT^Fezf2^ in 12 sessions from 6 mice each for all stimulations, whisker stimulation Kruskal-Wallis test (*γj(2) = 120.32, p = 7.4e-27)* with Tukey-Kramer post hoc multiple comparison test IT^PlxnD1 vs. PlxnD1^ vs. PTFezf2 vs. Fezf2 (*p=9.5e-10)*, IT^PlxnD1 vs. PlxnD1^ vs. IT/PT^PlxnD1 vs. Fezf2^ (*p=9.5e-10)*, PT^Fezf2 vs. Fezf2^ vs. IT/PT^PlxnD1 vs. Fezf2^ (*p=0.46),* orofacial stimulation Kruskal-Wallis test (*χ^2^(2) = 141.64, p = 1.7e-31)* with Tukey-Kramer post hoc multiple comparison test IT^PlxnD1 vs. PlxnD1^ _vs. PT_^Fezf2 vs. Fezf2^ (*p=9.5e-10)*, IT^PlxnD1 vs. PlxnD1^ vs. IT/PT^PlxnD1 vs. Fezf2^ (*p=9.5e-10)*, PT^Fezf2 vs. Fezf2^ vs. IT/PT^PlxnD1 vs. Fezf2^ (*p=0.03)*, visual stimulation Kruskal-Wallis test (*χ^2^(2) = 130.05, p = 5.7e-29)* with Tukey-Kramer post hoc multiple comparison test IT^PlxnD1 vs. PlxnD1^ vs. PT^Fezf2 vs. Fezf2^ (*p=9.5e-10)*, IT^PlxnD1 vs. PlxnD1^ vs. IT/PT^PlxnD1 vs. Fezf2^ (*p=9.5e-10),* PT^Fezf2 vs. Fezf2^ vs. IT/PT^PlxnD1 vs. Fezf2^ (*p=0.21)).* *p<0.05, **p<0.005, ***p<0.0005.

**Extended Data 5.**
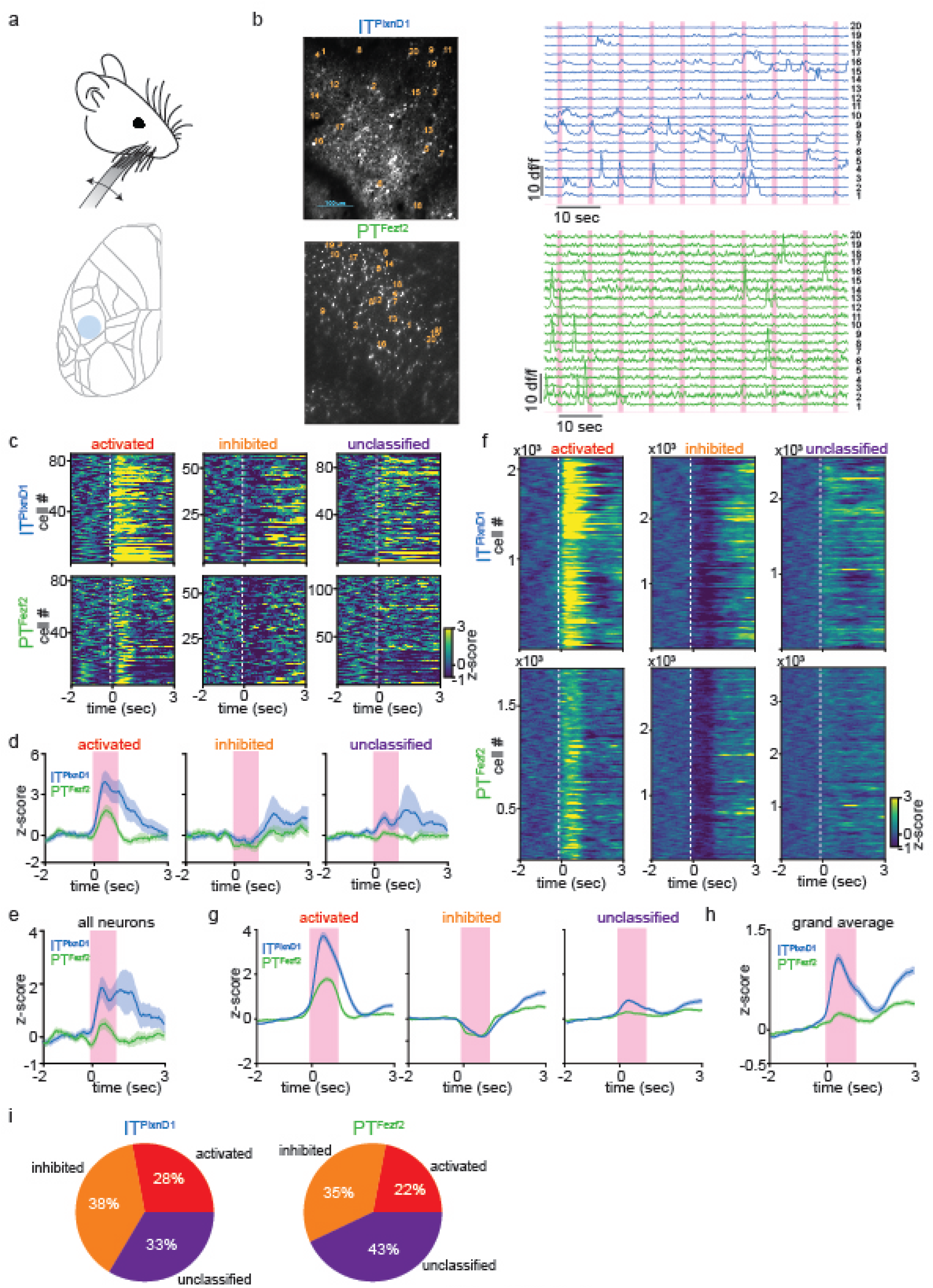
Calcium dynamics of IT^PlxnD1^ and PT^Fezf2^ at cellular resolution reflect widefield responses. a. Schematic of the whisker stimulation paradigm and the cortical location of two photon imaging (blue circle). b. Left: Example field of view (FOV) of single IT^PlxnD1^ cell bodies and apical dendrites of PT^Fezf2^ in the whisker barrel cortex. Right: Example traces from single IT^PlxnD1^ cell bodies and apical dendrites of PT^Fezf2^. Numbers indicate the corresponding location on the FOV. Magenta bars indicate whisker stimulation events. c. Heat map of average single neuron responses of IT^PlxnD1^ and PT^Fezf2^ classified into 3 groups based on their activity during whisker stimulation from the example FOV. d. Average responses across all IT^PlxnD1^ and PT^Fezf2^ neurons within each group from the example FOV. Magenta bars indicate duration of whisker stimulation. e. Average responses across all IT^PlxnD1^ and PT^Fezf2^ neurons from the example FOV. f. Heat map of average single neuron responses of IT^PlxnD1^ and PT^Fezf2^ classified based on their activity during whisker stimulation across all mice and sessions (IT^PlxnD1^ 42 FOV’s from n = 4 mice and PT^Fezf2^ 36 FOV’s from n = 3 mice). g. Average responses across all IT^PlxnD1^ and PT^Fezf2^ neurons within each group across all mice and sessions (IT^PlxnD1^ 42 FOV’s from n = 4 mice and PT^Fezf2^ 36 FOV’s from n = 3 mice). h. Average responses across all IT^PlxnD1^ and PT^Fezf2^ neurons from all mice and sessions combined (IT^PlxnD1^ 42 FOV’s from n = 4 mice and PT^Fezf2^ 36 FOV’s from n = 3 mice). i. Proportion of neurons in each group from IT^PlxnD1^ and PT^Fezf2^.

**Extended Data 6.**
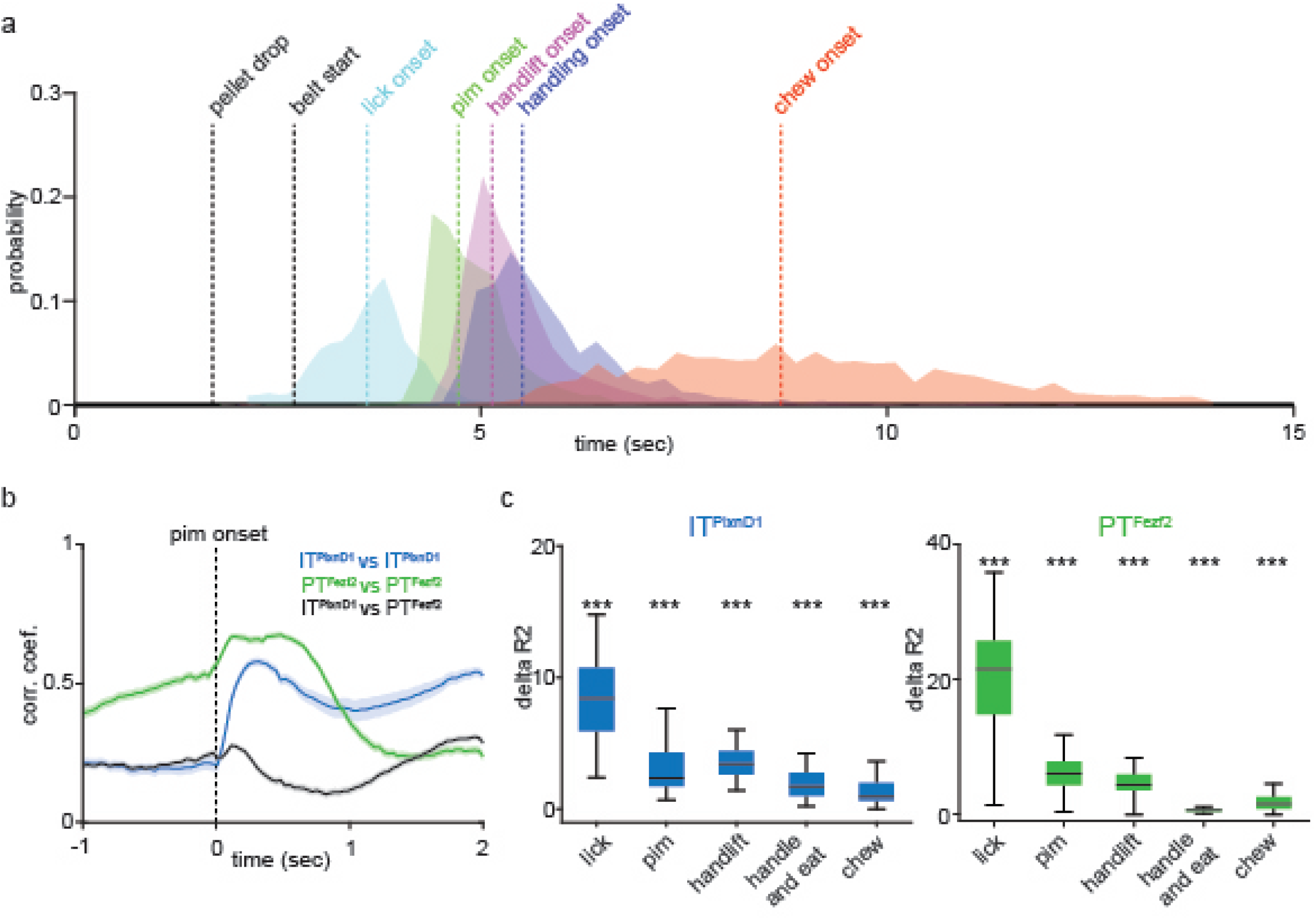
Correlation of activity maps during feeding behavior. a. Probability distribution of onset of lick, PIM, hand lift, first handling event and first chew event across all mice and sessions. Dashed lines indicate median times. b. Average Pearson’s correlation coefficient within IT^PlxnD1^ (blue), PT^Fezf2^ (green) and between IT^PlxnD1^ & PT^Fezf2^ mean activity maps at each time step before and after pellet-in-mouth onset (253 pairs within IT^PlxnD1^, 276 pairs within PT^Fezf2^ and 552 pairs between IT^PlxnD1^ & PT^Fezf2^ from 23 sessions in 6 IT^PlxnD1^ mice and 24 sessions in 5 PT^Fezf2^ mice). c. Distribution of the difference in variance explained between full and reduced model (delta R2, used in Fig2e) for each variable from IT^PlxnD1^ (top, 23 sessions from 6 mice) and PT^Fezf2^ (bottom, 24 sessions from 5 mice). ***p<0.0005.

**Extended Data 7.**
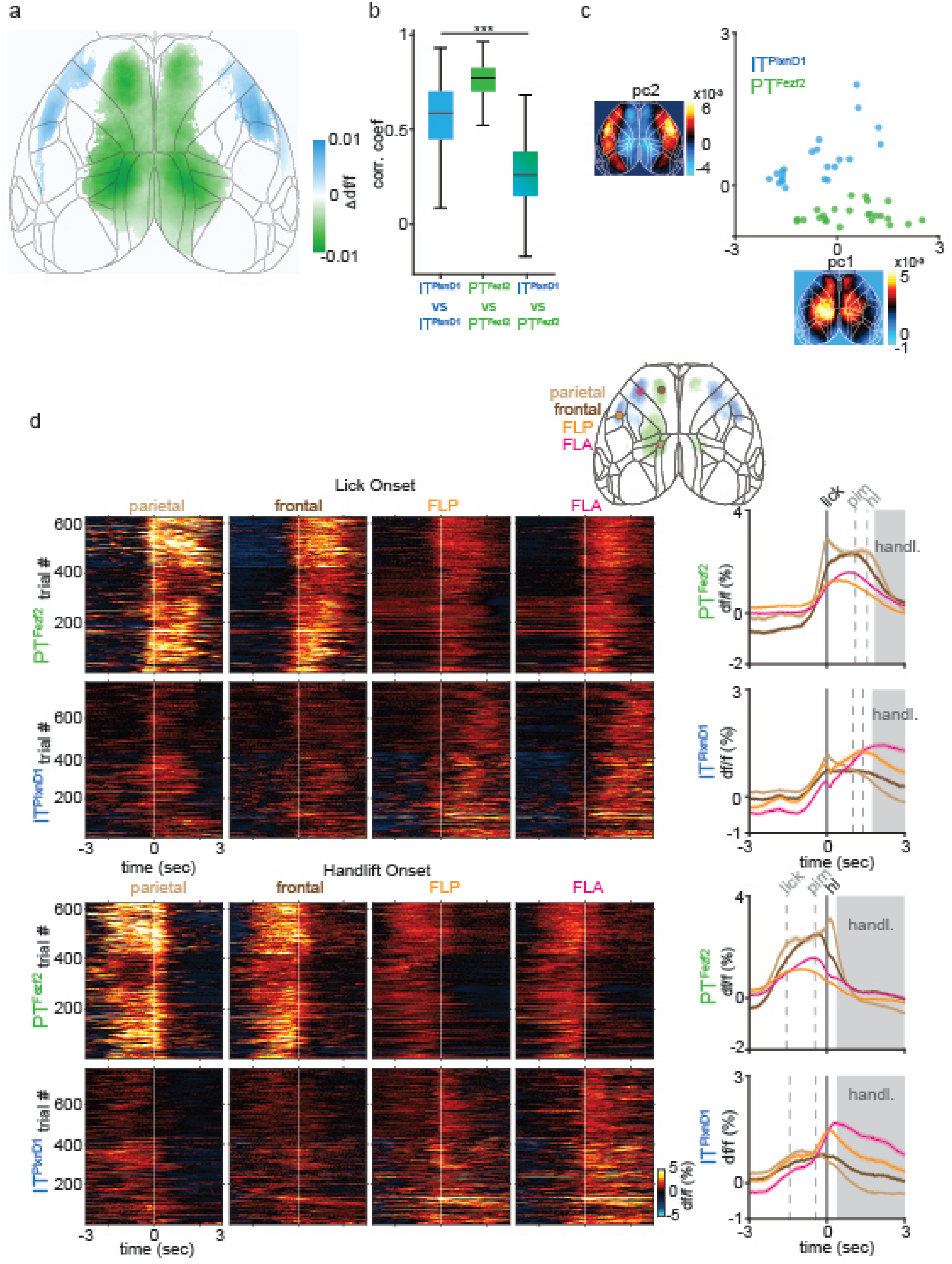
Temporal dynamics of IT^PlxnD1^ and PT^Fezf2^ within parietofrontal and frontolateral networks centered to lick and hand lift onset. a. Difference between IT^PlxnD1^ and PT^Fezf2^ mean activity map from Fig 3a,d. Only significantly different pixels are displayed (*two-sided Wilcoxon rank sum test with p-value adjusted by FDR = 0.05).* Blue pixels indicate values significantly larger in IT^PlxnD1^ compared to PT^Fezf2^ and vice versa for green pixels. b. Distribution of Pearson’s correlation coefficients within IT^PlxnD1^ (blue), PT^Fezf2^ (green) and between IT^PlxnD1^ & PT^Fezf2^ (blue green) mean feeding sequence activity maps (253 pairs within IT^PlxnD1^, 276 pairs within PT^Fezf2^ and 522 pairs between IT^PlxnD1^ & PT^Fezf2^, Kruskal-Wallis test (*χ^2^(2) = 681.83, p = 8.7e-149)* with Tukey-Kramer post hoc multiple comparison test IT^PlxnD1 vs. PlxnD1^ vs. PT^Fezf2 vs. Fezf2^ (*p=9.5e-10),* IT^PlxnD1 vs. PlxnD1^ vs. IT/PT^PlxnD1 vs. Fezf2^ (*p=9.5e-10),* PT^Fezf2 vs. Fezf2^ vs. IT/PT^PlxnD1 vs. Fezf2^ (*p=9.5e-10)).* c. Distribution of IT^PlxnD1^ (blue) and PT^Fezf2^ (green) mean feeding sequence activity maps projected to the subspace spanned by the top two principal components. d. Single trial heat maps and mean activity of PT^Fezf2^ and IT^PlxnD1^ from parietal, frontal, FLA and FLP centered to lick (top) and handlift onset (bottom, IT^PlxnD1^ - 23 sessions from 6 mice, PT^Fezf2^ - 24 sessions from 5 mice, shading around trace ±2 s.e.m). ***p<0.0005.

**Extended Data 8.**
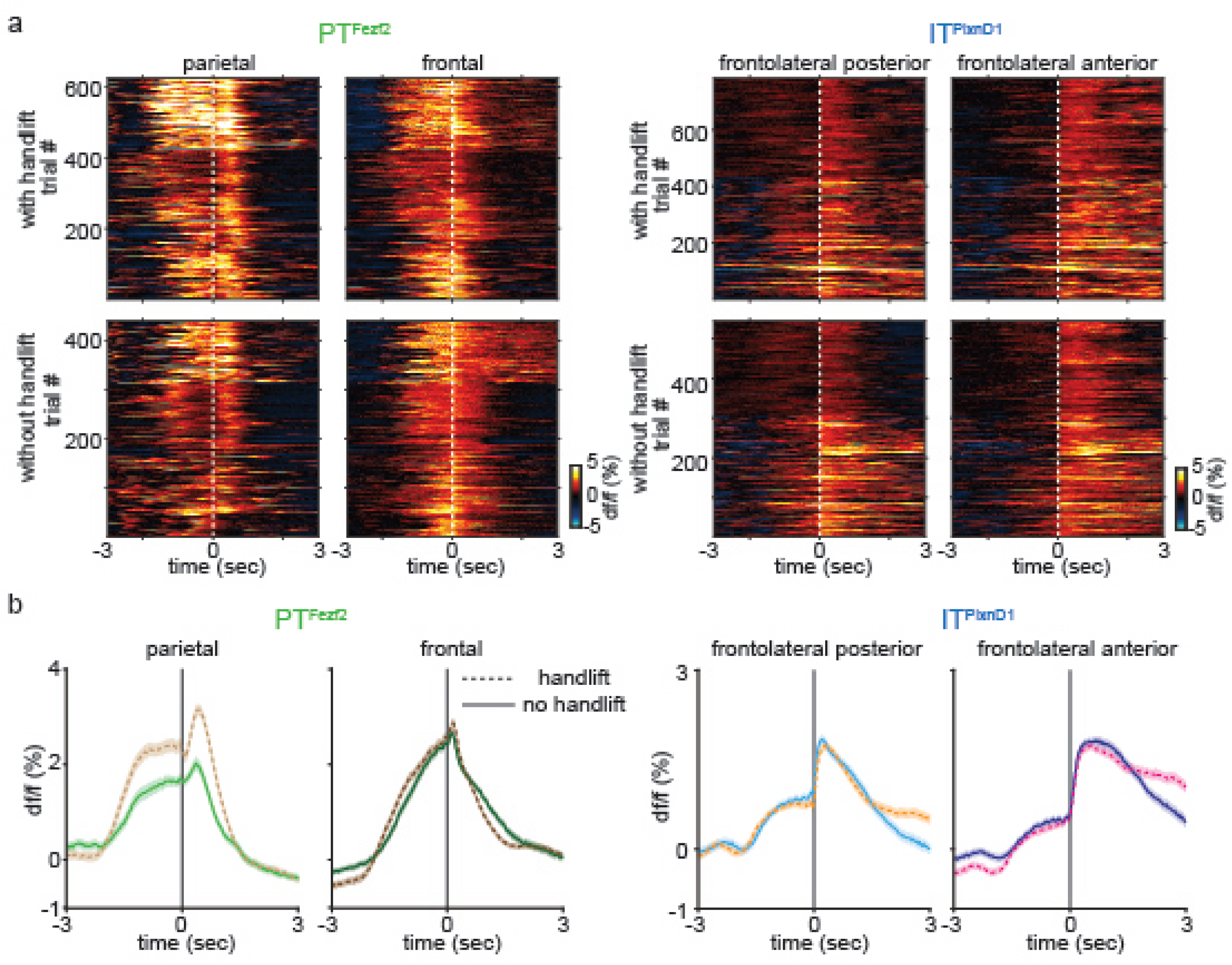
Temporal dynamics of IT^PlxnD1^ in frontolateral and PT^Fezf2^ within parietofrontal nodes during feeding with and without hand lift across mice. a. Single trial heat maps of PT^Fezf2^ activity centered to pellet in mouth onset from parietal and frontal node and IT^PlxnD1^ activity in FLP and FLA from eating with (top) and without hand lift (bottom) from all mice and sessions (With handlift : IT^PlxnD1^ - 23 sessions from 6 mice, PT^Fezf2^ - 24 sessions from 5 mice. Without hand lift: IT^PlxnD1^ - 15 sessions from 6 mice, PT^Fezf2^ - 13 sessions from 5 mice). b. Left: Mean PT^Fezf2^ activity aligned to pellet in mouth onset from parietal node with (light brown) and without (light green) hand lift, frontal node with (dark brown) and without (dark green) hand lift. Right: IT^PlxnD1^ activity aligned to PIM onset from FLP with (orange) and without (cyan) hand lift and FLA with (magenta) and without (dark blue) hand lift (shading around trace ±2 s.e.m).

**Extended Data 9.**
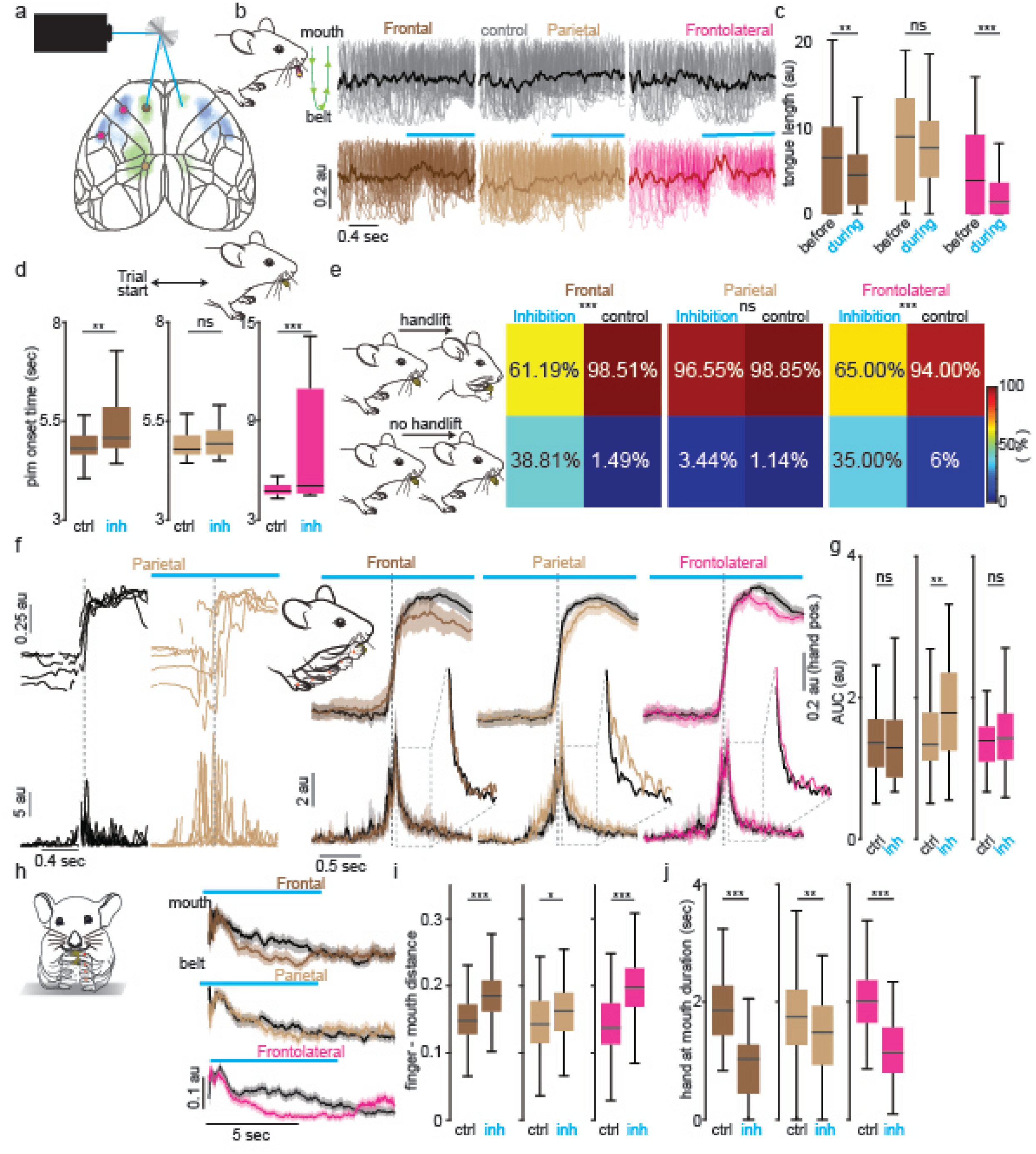
Inhibition of parietofrontal and frontolateral regions differentially disrupts sensorimotor components of feeding behavior. a. Schematic of optogenetic laser scanning setup. b. Single trial (translucent) and averaged (opaque) tongue trajectories centered to bilateral inhibition (bottom row) of frontal (dark brown), parietal (light brown) and frontolateral (magenta, pooled across FLA and FLP) nodes. Grey trajectories (top row) are from control trials preceding inhibition trials (n = 3 mice). Note that the trajectories evolve from top to bottom with the mouth at top and the pellet at bottom. An upward change in mean value indicates a decrease in tongue trajectory. Green schematic illustrates the direction of evolution of a single lick trajectory. c. Distribution of total tongue length 0.5 seconds before and after inhibition onset of frontal (dark brown), parietal (light brown) and frontolateral (magenta) nodes (paired two-sided Wilcoxon signed rank test, pooled from 3 mice across 7 sessions, 76 trials from frontal (*U=1522, z=3.22, p=0.0013),* 88 trials from parietal (*U=1919, z=1, p=0.31),* 89 trials from frontolateral (*U=1895, z=4.72, p=2.3e-6)*). d. Distribution of durations to pick pellet after trial start during control trials versus inhibition of frontal (dark brown), parietal (light brown), and frontolateral (magenta) nodes (two-sided Wilcoxon rank sum test, pooled from 3 mice across 7 sessions, 76 trials from frontal (*U=5393.5, z=3.32, p=9.13e-4),* 88 trials from parietal (*U=5919, z=0.67, p=0.5),* 89 trials from frontolateral (*U=6867, z=3.52, p=4.3e-4)).* e. Probability of hand lift events during bilateral inhibition of frontal, parietal frontolateral nodes compared to control (chi-Squared test, pooled from 3 mice across 7 sessions, 67 trials from frontal (*χ^2^=28.99, p=7.2e-8),* 87 trials from parietal (*χ^2^=1.02, p=0.31),* 100 trials from frontolateral (*χ^2^=25.80, p=3.78e-7)).* f. Left: 5 example hand lift trajectories while lifting hand to mouth from side view (top) during control (black) and inhibition of parietal node (light brown) and corresponding absolute velocities (bottom). Right: Mean vertical hand trajectory from side view (top, shading around trace ±2 s.e.m) and absolute velocity (bottom, shading around trace ±2 s.e.m) while lifting hand to mouth during control (black) and inhibition of frontal (dark brown), parietal (light brown) and frontolateral (magenta) nodes. Insets indicate zoomed in mean signals. Note the increase in velocity fluctuation during parietal inhibition. g. Distribution of the integral of absolute velocity signal for 1 sec post hand lift during control and inhibition of frontal (dark brown), parietal (light brown, p<0.005 Wilcoxon rank sum test) and frontolateral (magenta) nodes (two-sided Wilcoxon rank sum test, pooled from 3 mice across 7 sessions, 41 trials from frontal (*U=1769, z=0.62, p=0.53),* 83 trials from parietal (*U=5872, z=-3.42, p=6.3e-4),* 59 trials from frontolateral (*U=3356, z=-0.82, p=0.41)*). h. Mean normalized vertical trajectory of left finger from the front view after licking pellet into mouth during control trials (black, shading around trace 2 s.e.m) and inhibition of frontal (dark brown), parietal (light brown) and frontolateral (magenta) nodes. i. Distribution of mean normalized finger to mouth distance during control versus inhibition of frontal (dark brown), parietal (light brown) and frontolateral (magenta) nodes (twosided Wilcoxon rank sum test, pooled from 3 mice across 7 sessions, 58 trials from frontal (*U=2545, z=-4.68, p=2.87e-6),* 89 trials from parietal (*U=7222, z=-2.16, p=0.03),* 101 trials from frontolateral (*U=6925, z=-7.98, p=1.43e-15)).* j. Distribution of duration of hand held close to mouth during control versus inhibition of frontal (dark brown), parietal (light brown) and frontolateral (magenta) nodes (two-sided Wilcoxon rank sum test, pooled from 3 mice across 7 sessions, 58 trials from frontal (*U=4702, z=7.22, p=4e-13),* 89 trials from parietal (*U=9065.5, z=3.2, p=0.0014),* 101 trials from frontolateral (*U=13976, z=8.96, p=3e-19)*). *p<0.05, **p<0.005, ***p<0.0005.

**Extended Data 10.**
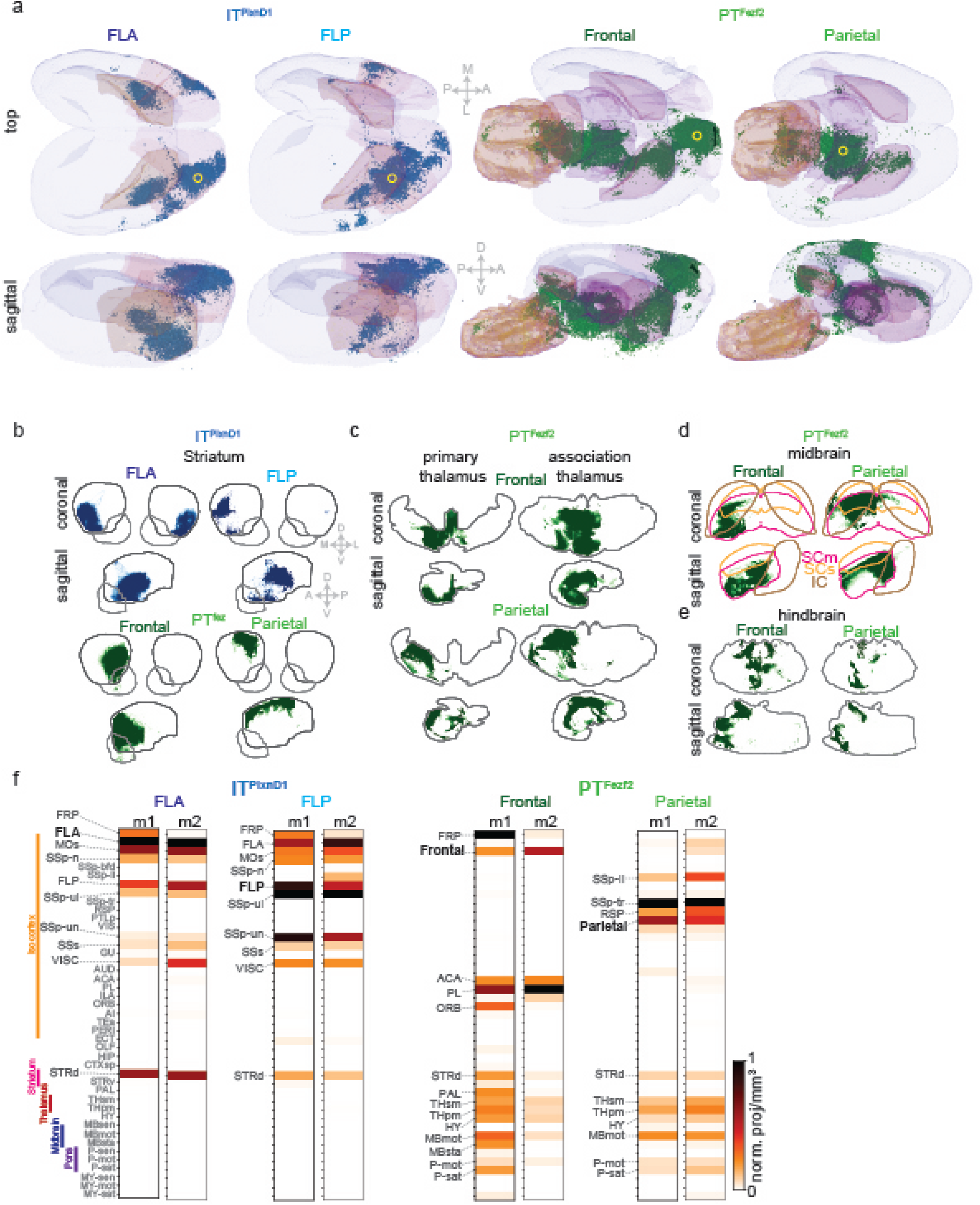
Axonal projection of IT^PlxnD1^ and PT^Fezf2^ in subcortical structures. a. Three dimensional rendering of axonal projections of IT^PlxnD1^ from FLA and FLP and PT^Fezf2^ from frontal and parietal node. Yellow circle indicates injection site. b. Spatial distribution of axonal projections of IT^PlxnD1^ from FLA and FLP (top) and PT^Fezf2^ parietal and frontal nodes (bottom) within the striatum projected onto the coronal and sagittal plane. c. Spatial distribution of axonal projections of PT^Fezf2^ from parietal and frontal nodes within the primary and association thalamus projected onto the coronal and sagittal plane. d. Spatial distribution of axonal projections of PT^Fezf2^ from parietal and frontal nodes within the motor Superior colliculus (SCm, magenta), sensory superior colliculus (SCs, yellow) and inferior colliculus (IC, brown) projected onto the coronal and sagittal plane. e. Spatial distribution of axonal projections of PT^Fezf2^ from parietal and frontal nodes (bottom) within the hindbrain projected onto the coronal and sagittal plane. f. Brain-wide volume and peak normalized projection intensity maps of IT^PlxnD1^ from FLA and FLP and PT^Fezf2^ from frontal and parietal nodes from two mice. Black font indicates injection site; larger gray font indicates regions with significant projections; smaller gray font indicates regions analyzed.

**Extended data 11.**
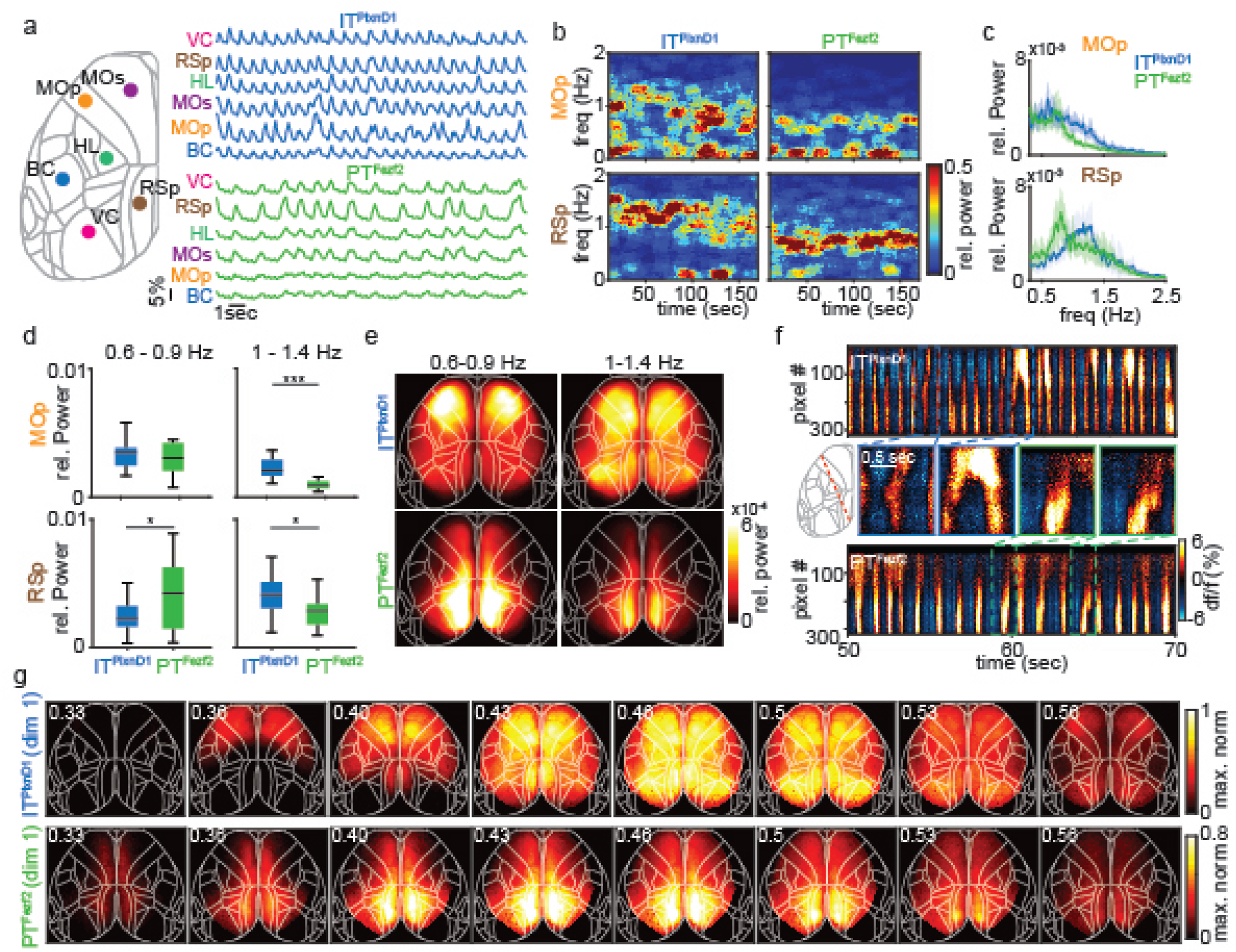
IT^PlxnD1^ and PT^Fezf2^ show distinct spatiotemporal dynamics and spectral properties under ketamine anesthesia. a. Example single trial traces of IT^PlxnD1^ (blue) and PT^Fezf2^ (green) activities from 6 different cortical areas during ketamine/xylazine anesthesia. Colors represent cortical areas as indicated (VC – primary visual Cortex, RSp – medial retrosplenial cortex, HL – primary hindlimb sensory cortex, MOs – secondary motor Cortex, MOp – primary motor cortex, BC – barrel cortex). b. Example spectrogram of IT^PlxnD1^ and PT^Fezf2^ activity from MOp and RSp of one mouse. c. Mean relative power spectral density of IT^PlxnD1^ (blue) and PT^Fezf2^ (green) activity from MOp and RSp (18 sessions across 6 mice each, shading around trace ±2 s.e.m). d. Distribution of average relative power within MOp and RSp of IT^PlxnD1^ and PT^Fezf2^ between 0.6-0.9 Hz and 1-1.4 Hz (18 sessions across 6 mice each, two-sided Wilcoxon rank sum test, 0.6-0.9 Hz MOp (*U=353, z=0.62, p=0.54),* RSp (*U=266, z=-2.1, p=0.03),* 1-1.4 Hz MOp (*U=482, z=4.7, p=2.6e-6),* RSp (*U=400, z=2.1, p=0.03)).* e. Spatial map of average relative power between 0.6-0.9 Hz and 1-1.4 Hz from IT^PlxnD1^ (top) and PT^Fezf2^ (bottom, 18 sessions across 6 mice each). f. Example space-time plots of the neural activity across a slice of the dorsal cortex (red dashed line) from IT^PlxnD1^ (top) and PT^Fezf2^ (bottom). Middle, zoomed-in activity from indicated top and bottom panels. g. Example activation sequence of the most dominant pattern (1^st^ dimension) identified by seqNMF from IT^PlxnD1^ (top) and PT^Fezf2^ (bottom). *p<0.05, **p<0.005, ***p<0.0005.

**Extended Data 12.**
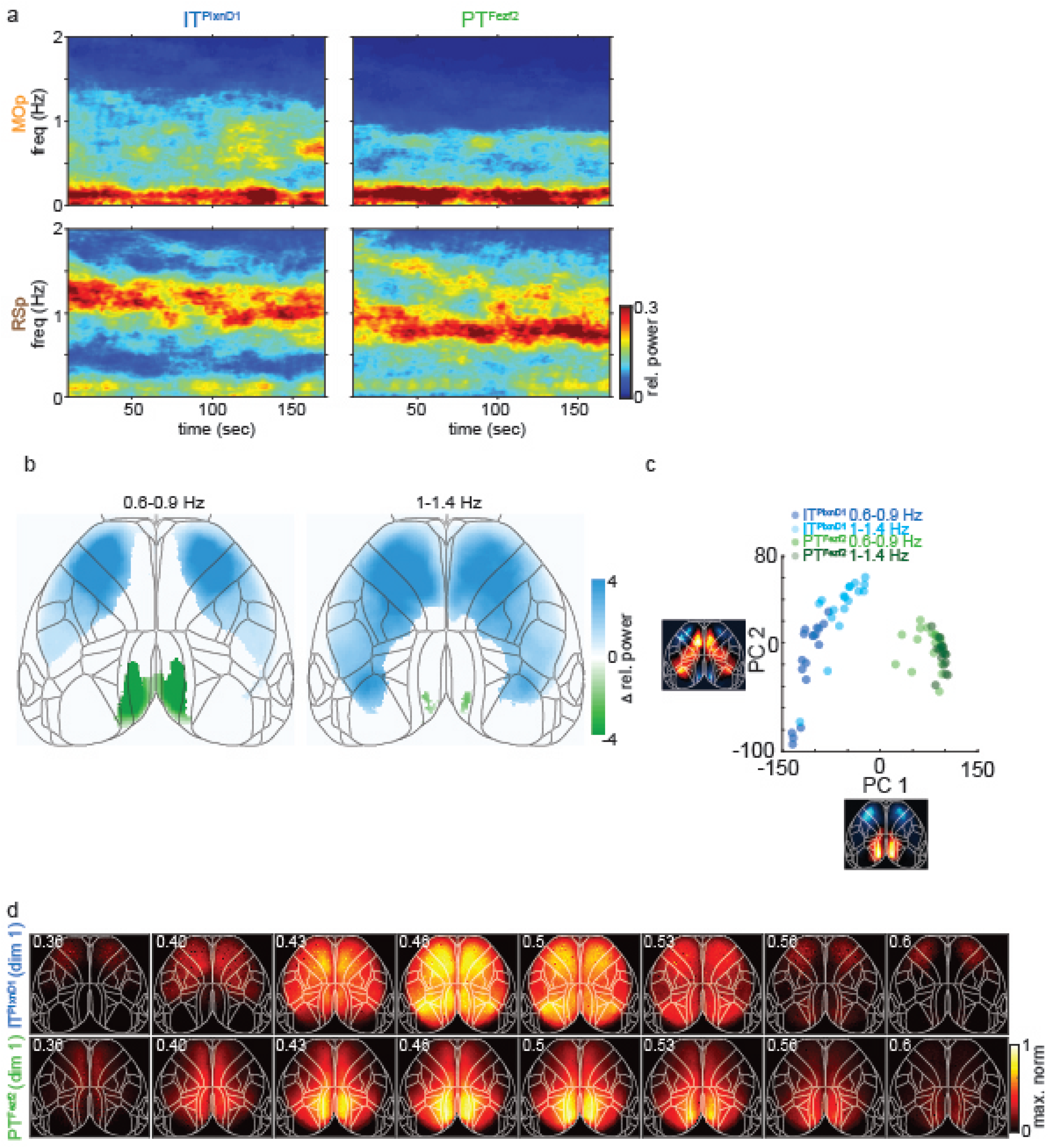
Spatiotemporal dynamics of IT^PlxnD1^ and PT^Fezf2^ under ketamine anesthesia across mice. a. Mean spectrogram of IT^PlxnD1^ and PT^Fezf2^ activity from MOp and RSp (18 sessions across 6 mice for each cell type). b. Difference between IT^PlxnD1^ and PT^Fezf2^ average relative power maps for each frequency bands (18 sessions from 6 mice for each cell type). Only significantly different pixels are displayed (*two-sided Wilcoxon rank sum test with p-value adjusted by FDR = 0.05).* Blue pixels indicate values significantly larger in IT^PlxnD1^ compared to PT^Fezf2^ and vice versa for green pixels. c. Distribution of IT^PlxnD1^ (blue) and PT^Fezf2^ (green) spatial power maps for each frequency band projected to the subspace spanned by the top two principal components. d. Activation sequence of the most dominant pattern (1^st^ dimension) identified by seqNMF from IT^PlxnD1^ (top) and PT^Fezf2^ (bottom) activity combined across mice and sessions.

**Extended Data 13.**
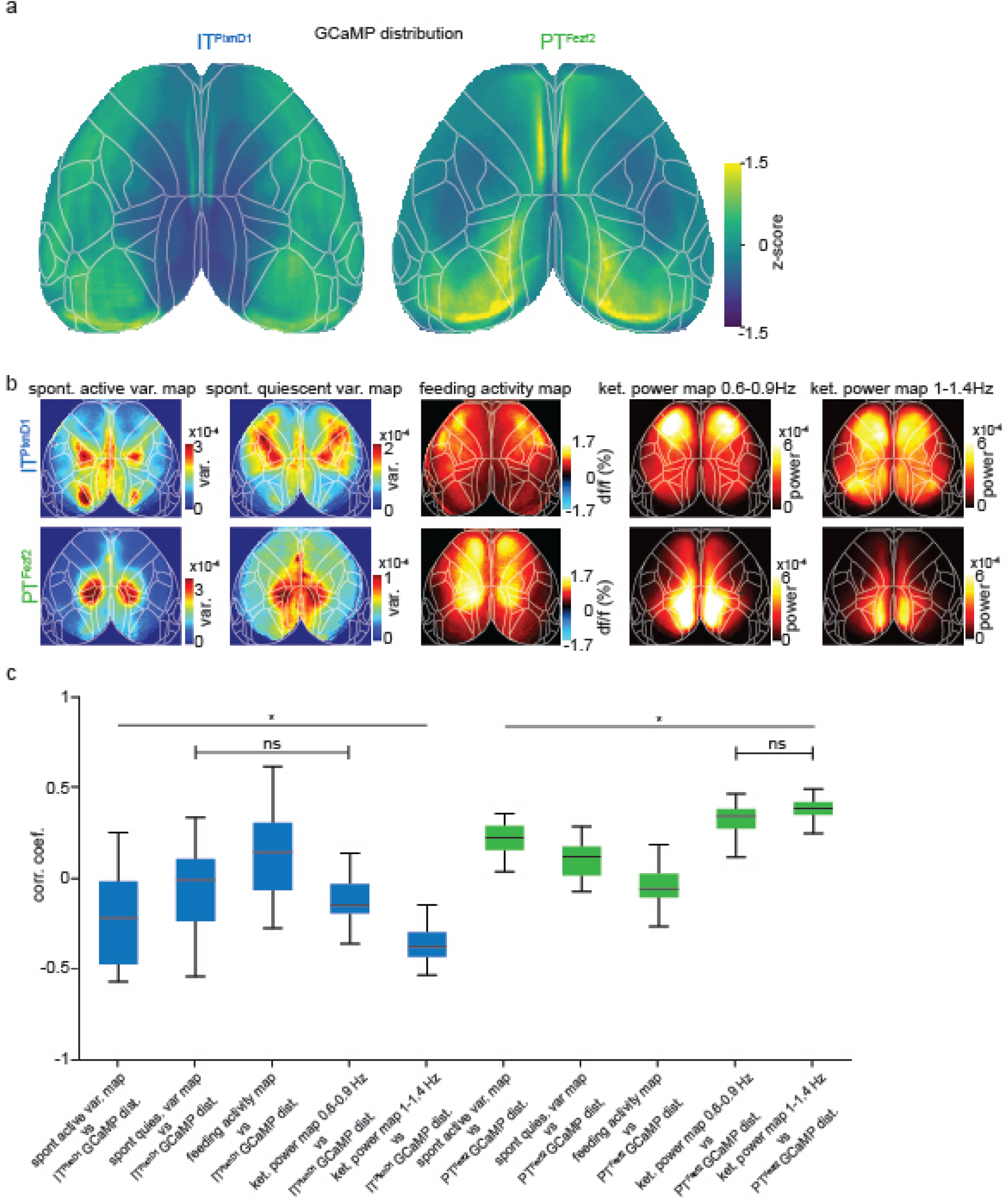
Correlation between IT^PlxnD1^ and PT^Fezf2^ GCaMP6f distribution and activity maps. a. Mean distribution of GCaMP6f density in IT^PlxnD1^ (left) and PT^Fezf2^ (right) neurons within cortex projected on to the dorsal cortex surface (n=4 mice for each cell type). b. Mean activity maps of IT^PlxnD1^ and PT^Fezf2^ across different brain states. From left to right: Mean active variance maps during spontaneous behavior (12 sessions from 6 mice for each cell type), mean quiescent variance maps during spontaneous behavior (12 sessions from 6 mice for each cell type), mean activity maps during feeding behavior(IT^PlxnD1^ : 23 sessions from 6 mice, PT^Fezf2^: 24 sessions from 5 mice), mean relative power maps for 0.6-0.9 Hz during ket./xyl. anesthesia (18 sessions from 6 mice for each cell type) and mean relative power maps for 1-1.4 Hz during ket./xyl. anesthesia (18 sessions from 6 mice for each cell type). c. Distribution of Pearson’s correlation coefficient between GCaMP6f cell density and each of the activity maps (from panel b) within the corresponding cell types (one way ANOVA (*F_9,658_=163.07, p=0)* with Tukey-Kramer post hoc multiple comparison test. All pairs of comparisons within each cell type are significantly different *(at* least with *p<0.05)* except for the two pairs indicated as non-significant (ns)).

## Video Captions

**Supplementary Video 1.**

Segment of single trial calcium activity across dorsal cortex during awake resting state from one example IT^PlxnD1^ mouse (with overlaid dorsal Allen map (CCF v3)) with temporally matched behavior video frames. Each frame is indicated as quiescent or active based on behavior variance crossing a predefined threshold.

**Supplementary Video 2.**

Segment of single trial calcium activity across dorsal cortex during awake resting state from one example PT^Fezf2^ mouse (with overlaid dorsal Allen map (CCF v3)) with temporally matched behavior video frames. Each frame is indicated as quiescent or active based on behavior variance crossing a predefined threshold.

**Supplementary Video 3.**

Example video of a single trial head fixed feeding behavior with body parts tracked using deeplabcut displaying both front and side view with simultaneous traces of the tracked body parts and behavior features.

**Supplementary Video 4.**

Single trial calcium dynamics across dorsal cortex during head fixed feeding paradigm from one example IT^PlxnD1^ mouse with temporally matched behavior video frames from front and side view.

**Supplementary Video 5.**

Single trial calcium dynamics across dorsal cortex during head fixed feeding paradigm from one example PT^Fezf2^ mouse with temporally matched behavior video frames from front and side view.

**Supplementary Video 6.**

Video demonstrating the effect on licking (tongue movement) after bilateral inhibition of frontal node in a PT^Fezf2^ mouse expressing GtACR1 (Right). Video on left shows behavior from the same mouse on the previous trial during the same period with no inhibition (control trial).

**Supplementary Video 7.**

Video demonstrating the effect on licking (tongue movement) after bilateral inhibition of frontal node in a IT^PlxnD1^ mouse expressing GtACR1 (Right). Video on left shows behavior from the same mouse on the previous trial during the same period with no inhibition (control trial).

**Supplementary Video 8.**

Video demonstrating the obstruction of handlift onset on bilateral inhibition of frontal node in a PT^Fezf2^ mouse expressing GtACR1 (Right). Video on left shows behavior from the same mouse on the previous trial during the same period with no inhibition (control trial).

**Supplementary Video 9.**

Video demonstrating the impairment in proper handling and manipulation of food pellet on bilateral inhibition of frontolateral anterior (FLA) node in a PT^Fezf2^ mouse expressing GtACR1 (Right). Video on left shows behavior from the same mouse on the previous trial during the same period with no inhibition (control trial).

**Supplementary Video 10.**

Video demonstrating the impairment in proper handling and manipulation of food pellet on bilateral inhibition of frontolateral anterior (FLA) node in a IT^PlxnD1^ mouse expressing GtACR1 (Right). Video on left shows behavior from the same mouse on the previous trial during the same period with no inhibition (control trial).

**Supplementary Video 11.**

Video of full brain STP imaged serial coronal sections with registered Allen map (CCF v3) overlaid from an IT^PlxnD1^ mouse injected with anterograde tracer in right frontolateral anterior node with its whole brain axonal projections.

**Supplementary Video 12.**

Video of full brain STP imaged serial coronal sections with registered Allen map (CCF v3) overlaid from an IT^PlxnD1^ mouse injected with anterograde tracer in right frontolateral posterior node with its whole brain axonal projections.

**Supplementary Video 13.**

Video of full brain STP imaged serial coronal sections with registered Allen map (CCF v3) overlaid from a PT^Fezf2^ mouse injected with anterograde tracer in right frontal node with its whole brain axonal projections.

**Supplementary Video 14.**

Video of full brain STP imaged serial coronal sections with registered Allen map (CCF v3) overlaid from a PT^Fezf2^ mouse injected with anterograde tracer in right parietal node with its whole brain axonal projections.

**Supplementary Video 15.**

Segment of single trial calcium activity across dorsal cortex during Ketamine/Xylazine anesthetized state from one example IT^PlxnD1^ mouse with overlaid dorsal Allen map (CCF v3).

**Supplementary Video 16.**

Segment of single trial calcium activity across dorsal cortex during Ketamine/Xylazine anesthetized state from one example PT^Fezf2^ mouse with overlaid dorsal Allen map (CCF v3).

